# High resolution nonlinear registration with simultaneous modelling of intensities

**DOI:** 10.1101/646802

**Authors:** Jesper L. R. Andersson, Mark Jenkinson, Stephen Smith

## Abstract

This paper describes and evaluates FMRIB’s nonlinear image registration tool (FNIRT), that is part of the FMRIB software library (FSL). It is a small deformation framework using sum of squared differences (SSD) as its cost function and Gauss-Newton for minimisation. The framework uses a joint shape and intensity model that attempts to explain the observed differences between two images in terms of having different shape and/or contrast, being differently affected by intensity bias-fields etc. Thus the estimation of the warps will be relatively unaffected by intensity differences that would otherwise violate the assumptions behind the SSD cost function. It uses a projection onto a manifold defined by a specified range of allowed Jacobian determinants to ensure that the warps are diffeomorphic. The utility of the model is demonstrated on a variety of simulated and experimental data with good results. FNIRT is also quantitatively evaluated using previously published datasets consisting of scans from multiple subjects, all with anatomically defined brain regions that are manually outlined. In this evaluation FNIRT performs well in comparison to previously published results with other registration algorithms.

## Introduction

Registering images of brains from different subjects is a necessary processing step for many types of analyses such as multi-subject fMRI studies, inter-group comparisons of tissue composition (e.g. VBM [1]), or measures derived from diffusion weighted MR (e.g. TBSS [2]). There is a family of algorithms that attempt to do this in the native “brain-space”, often referred to as “volumetric” (as opposed to “cortical” or surface based). These use different sets of transforms in native brain space between subjects, between a subject and a template in a standard space (atlas) or between longitudinal scans of the same subject. In order of increasingly “local” warps, the transforms can be divided into linear (affine) transforms, small deformation nonlinear transforms and large-deformation nonlinear transforms. The latter are typically framed in a way that is designed to guarantee diffeomorphic (topology preserving) warps.

These methods estimate the warps by minimising some scalar function of the parameters that define the warps. The function is designed to gauge how “different” the template is to the warped subject image, with the smaller the value the better the images “match”. One commonly used function is the sum-of-squared differences (SSD), which hinges on the assumption that the images are identical save for geometric differences. This assumption is frequently not fulfilled, e.g. because the two images are acquired with different modalities, have different contrasts due to differences in relative T_1_/T_2_-weighting or because the two images are differentially affected by *B*_1_ inhomogeneity (bias). This has lead to a multitude of different cost functions that all make slightly different assumptions and are designed to fare better when images differ in more respects than just geometry ([3], [4], [5], [6] and [7]). The question of which cost functions perform the best has been investigated in the context of linear (affine) registration (see [8] for an extensive review).

In contrast, the majority of methods for nonlinear registration of brain images use SSD ([9], [10], [11], [12], [13], [14], [15], [16], [17], [18]), though some have opted for mutual information ([19]). The latter makes less assumptions about the relationship between the images but may potentially be more susceptible to multiple local minima precisely for that reason. The mutual information approach addresses (as does for example correlation ratio [7]) issues of a global nonlinear relationship between the intensities in the two images, such as for example differences in degree of T1-weighting.

In our experience a more severe problem is local intensity changes, commonly caused by spatially varying sensitivity of the receive RF-coil [20]. With a birdcage coil, the dominant type of coil until recently, the intensity changes were of limited magnitude and consisted mainly of a small gradient in the *z*-direction. Recent developments in parallel imaging [21] have caused coil arrays [22] to become increasingly used, even for non-accelerated sequences. These consist of several small coils (32 coils are not uncommon and commercially available [23]) with highly focused and spatially non stationary sensitivity profiles. When this is combined with an accelerated acquisition and an appropriate reconstruction method (e.g. [24] and [25]) the resulting image will have a comparable scaling across the field of view (FOV) by virtue of the sensitivity of the coils being used in the reconstruction. However, the actual resulting sensitivity (summed over all coils) is typically considerably greater at the edges of the brain than in the centre. This means that when such data is combined with a non-accelerated acquisition followed by a simple sum-of-squares combination of the individual coil images the sensitivity (and hence intensities) will show a pronounced variation along all directions [26].

Furthermore the positioning of the subject inside the coil “helmet” has a substantial impact since the distance between the edge of the brain and the nearest coil will strongly affect the signal intensity. For example we frequently see a factor of 2-3 in intensity variation from posterior to anterior parts of the brain as a consequence of subjects being positioned below the centre of the coil-array in the superior-inferior direction. These intensity variations will invalidate the assumptions behind SSD, correlation ratio and mutual information alike and unless corrected will cause spurious warps in the regions affected by the inhomogeneities. Attempts have been made at addressing this by reformulating correlation ratio ([27], [28] and [29]) or mutual information ([30] and [31]) to render them “local” in that only a local region is considered when calculating it. This makes them less sensitive to intensity variations over scales that exceeds that of the, arbitrary, region over which the statistic is calculated. A different approach was taken by [32] who included an explicit bias field in a joint registration-segmentation framework where a set of prior tissue probabilities in standard space were warped so as to maximise the posterior probability of the segmentation of the registered image. A slight disadvantage of this method is that inter-subject or longitudinal registration will have to be performed by combining two individual transforms into standard space.

Present and future high field scanners will also have a problem with inhomogeneous deposition of RF-energy. This is not primarily an effect of inhomogeneous transmission, but rather of the object itself (the head) interacting with the *B*_1_-field. It leads to a variation in flip-angle across the FOV that means that the tissue contrast (as opposed to just the intensity) will differ across the FOV (see [33] for equations describing how tissue contrast depends on flip angle). This effect cannot be modelled by a multiplicative bias field (as is often done for RF-receive inhomogeneity) but would need a spatially varying nonlinear mapping of intensities.

The choice of cost function will also affect what methods one can use to search for the warps, i.e. to minimise the cost function. Any cost function that attempts to explain the “data” (the reference image in this context) in terms of a model and a normal distributed error lends itself to be minimised by the Gauss-Newton method [34] or one of its variants such as e.g. Levenberg-Marquardt. Both the SSD (where the model is simply the intensities of the image being registered) and the registration-segmentation model used by [32] are examples of cost functions fitting into this framework and where the Gauss-Newton method has been used (e.g. [11], [32], [18] and [35]). Other cost functions such as cross-correlation or mutual information do not fit into that framework and alternative methods have to be used.

There is a small conceptual divide between methods that are motivated from a parameter estimation perspective (e.g. [11] and [32]) and those that are formulated in terms of solving a set of partial differential equations PDE (e.g. [9], [10] and [12]) where these latter methods typically work by convolving a force field with some approximation to the Greens function of the differential operator of the PDE and integrating the resulting steps until the force field vanishes. This division is slightly artificial and models formulated as differential equations are in effect also parameter estimation problems that can benefit from methods traditionally used in that domain (e.g. [18]). An advantage of Gauss-Newton type minimisation is that it is often very efficient in terms of the number of iterations needed (even though each iteration can be quite expensive due to the need to calculate the Hessian) and that, through the Hessian, one can obtain information on the uncertainty of the estimated parameters (deformation fields).

In what follows we describe a framework where differences in shape and intensity are modelled simultaneously, thereby preventing the latter to influence or be interpreted as the former. This is done by introducing additional parameters that model intensity differences and by inferring on these at the same time as the parameters for shape. A series of intensity models will be introduced and demonstrated and the context in which each one is suitable is described. Within this framework pairs of images with differing contrast or differentially affected by bias can be registered by minimisation of the residual error, i.e. the SSD. This enables us to use Gauss-Newton type optimisation to find the parameters in an efficient way. We further suggest a multi-stage process where shape and intensity are jointly modelled up to certain warp resolution after which the intensity parameters are held constant and only the shape parameters are further refined at higher resolutions. This allows us to switch to a different optimisation strategy for the higher resolutions where the cost of calculating and storing the Hessian becomes prohibitive.

## Theory

### Notation

The registration is performed by warping one image (volume) to another. We will refer to the image we want to warp as the “input” image and the stationary image (the image we want to warp it to) as the “reference” image. The reference image will be denoted by **f** when referred to as a whole and can interchangeably be an *lmn ×* 1 column vector or an *l × m × n* volume. When referring to a single (scalar) value at a specified index (or voxel coordinate) **i** = [*i j k*] we will use *f*_**i**_. The collection of all indicies [*i j k*] of **f** is denoted by 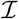 so an alternative notation for **f** would be 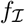. The warps (the nonlinear transformation) will be parametrised (the details of which can be found in Appendix S1) by a parameter vector **w**. The input image will be denoted by **g** when referred to as a whole in its original form. We use *g*_**i**_(**w**) to denote the value of **g** at the index **i** (as defined in the space of **f**) when warped by the parameters **w**. The notation **g**(**w**) is used to denote the values of **g** at all indicies 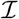 of the reference image **f** when applying the transform given by **w**. The functions *g*_**i**_(**w**) and **g**(**w**) are predicated on a specific interpolation model and they would for example be subtly different when spline interpolation is used compared to when using tri-linear interpolation.

### Registration as a nonlinear optimisation problem

Our optimisation problem can formally be described as

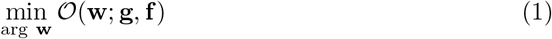

i.e. given **g** and **f**, find the set of parameters **w** that yield the minimum value for the cost function **𝒪**.

### Newton’s Method

Newton’s method for minimising **𝒪** is based on a first-order Taylor expansion of **𝒪** around some initial point **w**_0_, i.e.

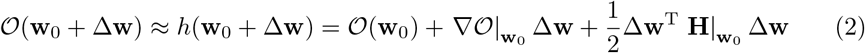

where ∇𝒪 is a column vector where the *i*th element is *∂*𝒪/*∂w*_*i*_ and **H** denotes the Hessian matrix where the *ij*th element is *∂*^2^𝒪/*∂w*_*i*_*∂w*_*j*_. The notation |**w**_0_ indicates that the adjacent entity was calculated at the point **w**_0_ in parameter space. Note that we have now defined a linear function *h*(**w**) that is an approximation of 𝒪(**w**) in some neighbourhood around **w**_0_. A necessary requisite for a minimum of 𝒪(**w**) is that ∇𝒪 = **0**. Hence, given some starting estimate **w**_0_ we wish to find the step ∆**w** that yields *∇𝒪*(**w**_0_ + ∆**w**) = **0**. However, there is no direct way to calculate that step so instead we use our approximate function and calculate the step ∆**w** that yields ∇*h*(**w**_0_ + ∆**w**) = **0**. From equation 2 we get

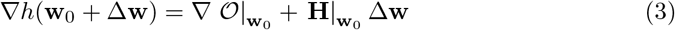

and by setting it to **0** and solving for ∆**w** we obtain

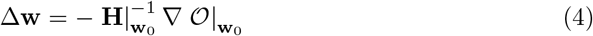

Since *h*(**w**) is an approximation of 𝒪(**w**) it is likely that ∇*h*(**w**_0_ + ∆**w**) ≠ ∇𝒪(**w**_0_ + ∆**w**), especially if ∆**w** is large (which implies that we are a long way away from **w**_0_ around which the approximation is valid). Therefore, to find the point where ∇𝒪 truly is **0** one uses an iterative scheme of updates to **w** where

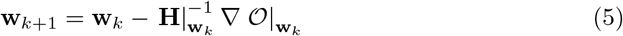

This is the update rule for Newton’s method for nonlinear optimisation, and the theoretical foundation for a multitude of derived methods.

From this point on we will simplify the notation by removing the |**w**_*i*_ whenever it is obvious from the context at which point the pertinent entity has been calculated.

### The Gauss-Newton approximation

There is a modification of Newton’s method, known as the Gauss-Newton method, that can be used when 𝒪(**w**) is of the form **o**^T^(**w**)**o**(**w**) where **o** is some *N*-dimensional vector-valued function. If one defines **J** as the matrix whose *ij*th entry is *∂o*_*i*_/*∂w*_*j*_ (where *o*_*i*_ is the *i*th element of **o** and *w*_*j*_ the *j*th element of **w**) one can write ∇𝒪 = 2**J**^T^**o** and **H** = 2**J**^T^**J** + **R**. The Gauss-Newton method is based on approximating **H** with 2**J**^T^**J**, i.e. ignoring **R**. An explanation for why this is feasible can be found in [34].

### Levenberg-Marquardt for increased robustness

Even though the Gauss-Newton approximation tends to be more robust than Newton’s method there is still a risk that a Gauss-Newton step can lead to a point with a higher value for 𝒪(**w**). A solution to this is to modify **J**^T^**J** so that is becomes more diagonal dominant whenever this happens. One such scheme is known as the Levenberg-Marquardt algorithm, which adaptively varies its behaviour between that of gradient-descent and Gauss-Newton [34].

### Alternative methods that do not need H

Even though it is typically, and definitely in the case of registration, faster to calculate **J**^T^**J** than the full **H** it is still a substantial task when the number of parameters (the length of the vector **w**) is large. Methods have therefore been developed that are also based on Eq 2 but which avoid calculating, and in some cases representing, **H**. The Conjugate Gradient methods find a set of direction **p**_*i*_of the form

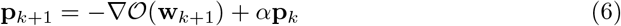

where a line-search is performed along each direction **p**_*i*_.

The Scaled Conjugate Gradient (SCG) method [36] is a further development that calculates also a step-length along **p**_*k*_, eliminating the need for a line-search. Analogously to the Levenberg-Marquardt algorithm it adjusts the step-length based on the outcome of the previous iteration.

### When is it an advantage to actually know/calculate H?

There exist problems for which knowing (calculating) **H** offers little advantage over and above just knowing ∇𝒪, and there are other problems for which it is crucial and where convergence would take a very long time without it. But which are which? The Hessian tells us for how long the derivative is “valid”. Let us say we have some function 𝒪(**w**) and let us say that the *i*th element of the gradient ∇𝒪 is large. That means that when taking our next step it should have a large component of (the negation of) the *i*th element, and we would consequently expect 𝒪 to decrease a lot. But what if **H**_*ii*_ is also large, how would that influence what direction we want to go in? That would mean that even though 𝒪 changes rapidly as we move along the *i*th direction it will soon stop changing so rapidly, and we may therefore not want to go very far. This consideration is executed by the multiplication with a small value for [**H**^−1^]_*ii*_ (assuming a diagonally dominant **H**). Likewise, if **H**_*ij*_ is large one would opt to go less distance in either the *i*th or the *j*th direction than one would otherwise do.

When using a method without explicit knowledge of **H** one will instead perform consecutive line minimisations along some set of directions in parameter space. The execution time will depend on the number of line minimisations one needs to perform. This will in turn depend on one’s choice of directions (i.e. minimisation algorithm), but more fundamentally it will also depend on the structure of the underlying (unknown) Hessian. If the Hessian is spherical the gradient will point towards the minimum and a single line minimisation would take one there. The other extreme would be when the Hessian is very complex with highly heterogeneous values both on and off the diagonal. In the worst case one may then need to perform as many line minimisations as there are directions (parameters), i.e. in the case of nonlinear registration several thousands to millions which would in practice not be realistic.

In between these extremes are cases where the Hessian divides up the parameter-space into a number of subspaces each of which is spherical, and one can then (at best) get away with performing as many line minimisations as there are such sub-spaces.

In the context of nonlinear registration the form of **H** depends on how the displacement fields are modelled. When using a basis-set with global support (such as e.g. the discrete cosine transform [11]) **H** will have a very complex structure and attempting to minimise a cost function without knowledge of **H** would result in a very large number of line minimisations and hence long execution time. In contrast a basis-set with local support (e.g. B-splines or “free-form” displacements) yields a diagonal dominant Hessian with a limited number of non-zero off-diagonal elements, which means that it is feasible to use a minimisation method without explicit use of **H**.

### The sum of squared differences cost function

The sum of squared differences (SSD) cost function is predicated on the model

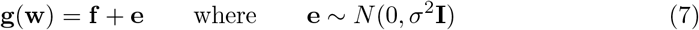

which gives a likelihood that is maximised when

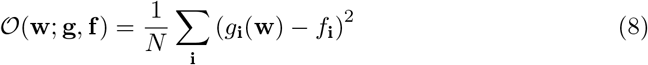

is minimised. To minimise this with the Gauss-Newton method we need to be able to calculate ∇𝒪 at any point **w** in parameter space. If we assume that the displacements are modelled by some basis-set parametrised by **w** then ∇𝒪 is a vector of the same size as **w** where the *j*th element is given by

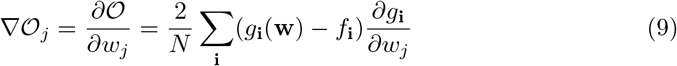

In vector-matrix notation ∇𝒪 can be written as

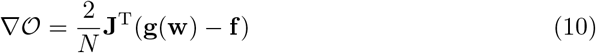

where **J** is the Jacobian matrix of **g**(**w**), i.e. the matrix whose *ij*th element is *∂g*_*i*_/*∂w*_*j*_. Using the same notation we can express the Gauss-Newton approximation of the Hessian as

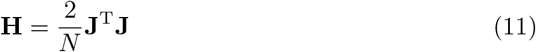

The details of how to calculate **J** depends on the basis-set that is used to represent the warps and will be relegated to Appendix S1.

### Simultaneous modelling of shape and intensities

The SSD has the advantage that it makes it straightforward to calculate ∇𝒪 and **H**, but it has the disadvantage that the underlying assumption that *f* and *g* are identical save for some difference in shape is almost always wrong. The assumptions of other cost functions, like e.g. Mutual Information, on the other hand are much more likely to be fulfilled. The disadvantage of those is that the elements of ∇𝒪 and **H** will have to be calculated numerically making the former very time consuming and the latter in practice infeasible.

The main contribution of this paper is to suggest a cost function that retains the advantages of SSD (straightforward analytical calculation of ∇𝒪 and **H**) while relaxing the very strict assumption that **g**(**w**) = **f** + **e** where **e** *~ N*(0, Σ). We do this by introducing a function *E*_**i**_(**b**; *f*_**i**_) that yields the “estimated” intensity at the location **i** given some set of parameters **b**. The simplest example of such a model would be where **b** is a scalar global scaling factor in which case *E*_**i**_(**b**; *f*_**i**_) = *bf*_**i**_. We have defined and implemented a hierarchy of such intensity models where the “lower level” models are special cases of those higher on the hierarchy. These are:

### None

As indicated by the name this implies no intensity modelling at all, i.e.

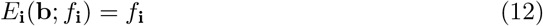

This may be useful for registration of “quantitative” images such as e.g. Fractional Anisotropy (FA).

### Global linear scaling

Here

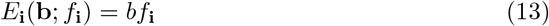

For this to be useful **f** and **g** should have been acquired with identical (or very similar) sequences (resulting in very similar contrasts) and both be unaffected by any RF inhomogeneity.

### Local linear scaling

For this model

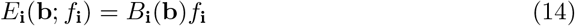

where *B*_**i**_(**b**) refers to the value of some scalar field **B**(**b**) at the location given by **i**. The mapping **b** *→* **B** is in principle arbitrary but will in our case be a set of continuous basis-functions with **b** as the coefficients. This is useful when **f** and **g** have similar contrast but are affected by RF inhomogeneity in different ways.

### Global nonlinear scaling

A polynomial of order *n −* 1 is used to model the intensities so that

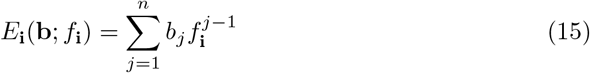

This can, within reason, model differences in amount of T1/T2-weighting between the two images.

### Global nonlinear scaling with bias field

This is a combination of the two preceding models where

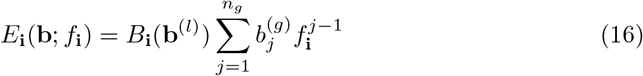

where **b** = [**b**^(*l*)^ **b**^(*g*)^], i.e. has been divided into a local and a global part. This models both differences in T1/T2-weighting and a bias field caused by differences in RF inhomogeneity.

### Local nonlinear scaling

This model encompasses all the previous ones and models *E*_**i**_(**b**; *f*_**i**_) as

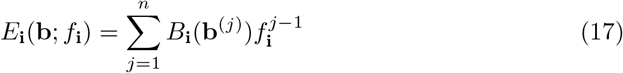

where **b** has now been divided into *n* sub-vectors, i.e. **b** = [**b**^(1)^ **b**^(2)^ … **b**^(*n*)^], each defining a field **B** of polynomial coefficients. Each coefficient will have a unique value at every voxel, but a given coefficient (say for example the second order coefficient) varies smoothly over space. Hence this model allows different nonlinear mappings (between the intensities in **f** and **g**) for different voxels.

It may seem superfluous with more than one model given that the model given by Eq 17 encompasses the other ones. However, the higher level models have more parameters which translates into longer execution times and potentially more local minima for a minimisation algorithm to fall into, i.e. a true geometric/anatomical difference my be modelled as an intensity difference if the intensity model is “too permissive”. As we will see later the opposite is also true, i.e. if the intensity model fails to explain the “true” intensity differences it will be modelled as geometric differences leading to non-sensical warps.

It should further be noted that even though the **b** *→* **B** mapping (in Eqs 14, 16 and 17) is in principle arbitrary it has to be such that the resulting field **B** is smooth compared to the resolution of (the image) **f**.

### ∇𝒪 and H when adding intensity modelling

When one simultaneously models shape and intensity there are two sets of parameters; **w** which parametrises the geometric warps and **b** which models intensity differences. Therefore ∇𝒪 changes so that

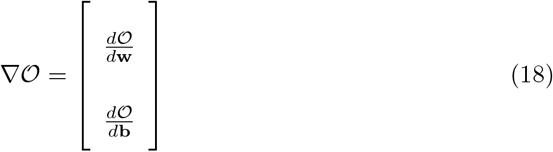

where *d*𝒪/*d***w** is a columnvector whose *i*th element is *∂*𝒪/*∂w*_*i*_ and *d*𝒪/*d***b** is one whose *i*th element is *∂*𝒪/*∂b*_*i*_ and **H** changes so that

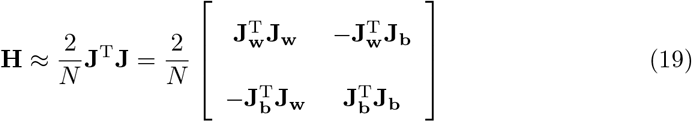

where **J_w_** is the Jacobian of the **w** *→* **g** mapping **g**(**w**) and **J_b_** is the Jacobian of the **b** *→* **E** mapping **E**(**b**; **f**). In Appendix S1 we derive expressions for ∇𝒪 and **H** for the different intensity models.

It should also be noted that the magnitude of the elements of *d*𝒪/*d***w** and *d*𝒪/*d***p** are *very* different, and hence 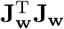, 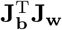 and 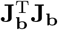 are of different orders of magnitude. That means that this is an example of when it is crucial to know, and use, **H** for the minimisation.

### High resolution warping

A disadvantage of explicitly using and representing **H** is the memory requirements. Let us say we have an image volume with an 180 × 220 × 180mm FOV and that we want to estimate warps with a resolution of 4mm. That means that **w** alone (i.e. ignoring **b**) consists of ~ 355500 elements which means that **H** is a 355500 × 355500 matrix with potentially ~ 6.3 ⋅ 10^10^ unique elements requiring ~ 500GB if one stores them with double precision. This number can be decreased considerably by choosing a suitable representation of the warps such that one can take advantage of the ensuing sparsity and symmetries, but it is nevertheless the case that calculating and representing **H** is the main difficulty when estimating high resolution warps. On the other hand, as indicated in the previous section, we need to know **H** when performing the joint warp and intensity modelling.

However, it is likely that the intensity parameters (**b**) can be estimated, using a Gauss-Newton-based algorithm, with “sufficient accuracy” together with a low/medium-resolution set of warps (**w**). These (**b**) could then be used as constants in a subsequent estimation of **w** at higher resolution. The elements of **w** are now the only parameters and, with a suitable representation of the warps, this means that **H** has a structure that is “close to” spherical and that it is feasible to use e.g. the SCG algorithm described above to find the warps.

### Regularisation

Given a large enough set of parameters **w** (and **b**) it is in principle possible to obtain an exact match between **g**(**w**) and **f**, but that might mean that the coordinate transform makes little sense with nearby points in the original space mapping to widely separated points in the target space, and vice versa. Additionally there may be a large set of different transforms that all yield similar values for 𝒪(**w**) and one needs a way of deciding which of those is the “best”. This is obtained by combining the SSD cost function with a regularisation term that is a scalar function *S*(**w**), which has a higher value for “less likely” fields, i.e. one that penalises non-smooth warp fields. This is equivalent to using a Normal prior on **w** so that **w** ~ *N* (0, *λ*^−1^**C**^−1^) where *S*(**w**) = *λ***w**^T^**Cw**. Hence, the cost function becomes

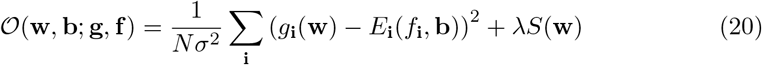

where *σ*^2^ is the sample variance and *λ* is an arbitrary factor that determines the relative weight of the regularisation. In the present paper *σ*^2^ could have been lumped into *λ* since it is an empirically determined “fudge-factor”. It is, however, still convenient to keep as a separate factor since it facilitates empirically determined values for *λ* that are valid over a range of different *σ*^2^ (i.e. different SNR). It also means that at earlier iterations when the estimated *σ*^2^ is large the regularisation is given more weight which helps avoiding local minima.

In addition, the intensity models described by Eqs 14, 16 and 17 all model maps representing either *B*_1−_ or *B*_1+_ fields, both of which are smooth and relatively slowly changing. Including this yields

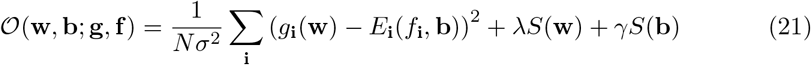

where *S*(**b**) is a function of the same form as *S*(**w**) operating on one (Eqs 14 or 16) or more (Eq 17) fields. As implemented in FNIRT *S*(**w**) and *S*(**b**) can be either “membrane energy” or “bending energy” (see e.g. [11]), though all experiments in this paper was performed using “bending energy”.

### Diffeomorphic mapping

#### What is a diffeomorphic mapping

Let us say we have two spaces *u* and *v* and that there is an 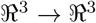 mapping from *u* to *v*. A diffeomorphic mapping is one which is “one-to-one”, meaning that each point in *u* maps onto a unique point in *v*, and “onto” which means that for every point in *v* there is some point in *u* that maps to it. Failure of the first condition is signalled by the Jacobian (the Determinant of the Jacobian matrix) of the mapping being *≤* 0 somewhere in the domain of *u*, while a failure of the second is more difficult to detect (and is typically ignored). The regularisation helps, but does not ensure that this condition is fulfilled. By giving the regularisation a large weight one can be almost guaranteed a diffeomorphic mapping, but that will, on the other hand, mean that the warped image (**g**(**w**)) will still be quite dissimilar to **f**. We will have what is commonly known as a “small deformation transform”.

#### Projection onto a diffeomorphic surface

There are methods that (almost) guarantee diffeomorphic mappings by virtue of the way in which they combine many small (diffeomorphic) deformations (see e.g. [15], [16], [18] or [35]). In the present paper we opted for a different solution, as proposed by [37]. We allow the field to become non-diffeomorphic for a limited number of iterations after which it is projected onto the “closest” diffeomorphic field. The partial derivatives of the fields (i.e. *∂φ*_*x*_/*∂x*, *∂φ*_*x*_/∂*y*,…, *∂φ*_*z*_/*∂z*) are explicitly manipulated/changed to ensure that the Jacobian determinant lies within a defined range. Following that, there are three derivative-maps for each component-field, that could each be used to calculate that component-field (i.e. *φ*_*x*_ could be calculated from either of *∂φ*_*x*_/*∂x*, *∂φ*_*x*_/*∂y* or *∂φ*_*x*_/*∂z*), but because of the changes that have been performed it is possible that the resulting fields are inconsistent (i.e. integrating *∂φ*_*x*_/*∂x* in the *x*-direction may yield a different result from integrating *∂φ*_*x*_/*∂y* in the *y*-direction. This potential inconsistency is resolved by Fourier-transforming the derivative maps, performing the projection onto the set of consistent gradient images in Fourier-space and transforming back into image-space.

## Materials and methods

### Implementation

The cost functions described above have been implemented in C++ in an application called FNIRT as part of the FSL software package (http://www.fmrib.ox.ac.uk/fsl). It has a command-line interface and is, in addition, an integral part of FEAT, FSL-VBM, TBSS and FDT. Some implementation details are given in Appendix S1. Here we will limit the description to say that there is a choice between the Levenberg-Marquardt (LM) and the Scaled Conjugate Gradient (SCG) methods for minimisation. The former explicitly calculates and uses the Hessian (**J**^T^**J**) and is hence quite memory hungry for high resolution (many elements in **w**). The latter does not explicitly use the Hessian and therefore needs less memory. The flip-side is that the SCG method cannot be used for simultaneous estimation of warp (**w**) and intensity (**b**) parameters since the implicit approximation of **H** is too poor in that case. FNIRT is able to take **w** and **b** as input, providing an initial guess for both. Therefore our strategy for achieving high resolution warps is to run FNIRT to a medium resolution (8–10mm knot spacing) with simultaneous estimation of intensity parameters **b**. A second run is then performed using the warp and intensity parameters from the first run as input, refining the former by going to a higher warp resolution but keeping the latter fixed.

## EXPERIMENTS

### Demonstration of intensity models

A set of simple simulations was performed to demonstrate the utility of the different intensity models. The “phantom” consisted of a set of geometric shapes: spheres, circular disks and a circular bowl. These were combined in such a way that when cut at a certain level in the *z*-direction it would resemble either a smiley or grumpy face when viewed in the *xy*-plane. These shapes were “imaged” using a 64 × 64 × 64 matrix with a 1 × 1 × 1mm voxel size. Intensities were assigned to the face, eyes and mouth to mimic those of white and grey matter and CSF respectively. To ensure that we assigned “realistic” intensities to the different tissue types, and also to introduce realistic intensity differences due to sequence differences and variable flip-angle, we used the signal equations for an MPRAGE sequence as outlined in [33]. For all simulations we used a relative proton density of 0.75/0.65/1.0 for grey matter/white matter/CSF. For the smiley face we used a *T*_*R*_ of 9.2ms, *T*_*I*_ of 0.9s, a delay time (*T*_*D*_) of 0.56s, 80 partitions per acquisition block (*n*) and a flip-angle (*α*) of 8 degrees, these being parameters that are commonly used at our lab, and *T*_1_ values of 1.61/0.84/4.3s for grey matter/white matter/CSF (relevant for 3T according to [38]). The same parameters were used for one of the grumpy faces to provide an image for registration with identical intensities. For the remaining grumpy faces we used 10ms/1s/0.5s/176/12° for *T*_*R*_/*T*_*I*_ /*T*_*D*_/*n*/*α* respectively and *T*_1_ values of 1.2/0.65/4.3s for grey matter/white matter/CSF (relevant for 1.5T according to [38]). The second grumpy face was created using these parameters with homogeneous *B*_1−_(receive *B*_1_ field) and *B*_1+_ (transmit *B*_1_ field) fields. An inhomogeneous field was created as a quadratic spline field with a knot spacing of 20mm where selected coefficients were set to 0.5 while the remaining ones were kept a 1. Quadratic splines were used so that the field should not be created with exactly the same basis as we use to model it (though it is admittedly very similar). This field was applied as a *B*_1−_ field to the third grumpy face (by element-wise multiplication) and as a *B*_1+_ field to the fourth grumpy face (by letting it modulate *α* in the signal equations in [33]). In all, this gives one smiley face to use as the target for all registrations and four grumpy faces whose intensities are successively more different from that of the smiley face.

The images generated by the simulations are shown in Fig 1. For display purposes they have been scaled such that the intensity in white matter (the face) is unity in all images. It can be noted that, as intended, the middle image has a different contrast from the two leftmost ones, mainly noticeable as a higher intensity in gray matter (the eyes). The second image from the right suffers from an additional inhomogeneous receive *B*_1_ field (*B*_1−_) leading to a local, multiplicative, reduction in intensities. The rightmost image was simulated with an inhomogeneous transmit *B*_1_ field (*B*_1+_) yielding a spatially varying flip-angle leading to a spatially varying contrast. This can be appreciated from the visually detectable increased contrast of the left eye to the surrounding face in the rightmost image compared to the one on its left.

**Fig 1.**
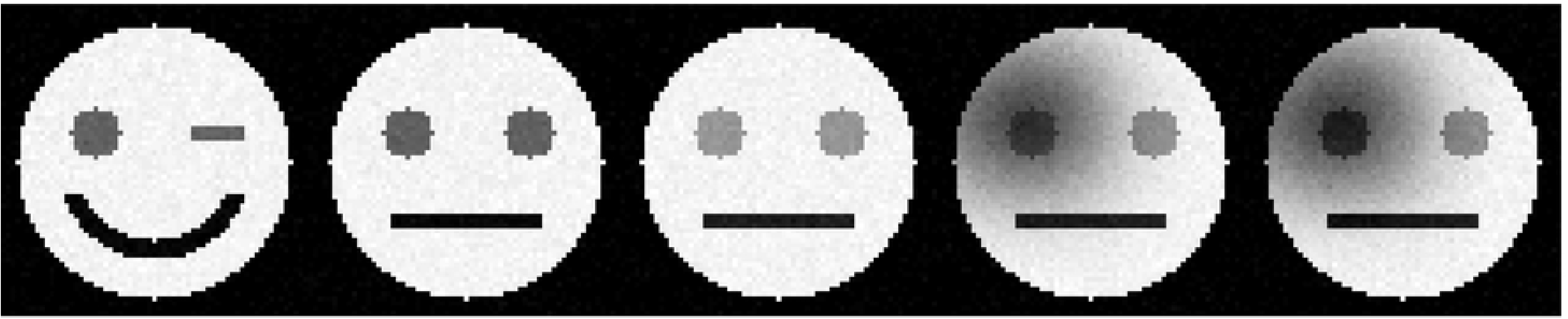
Phantoms simulated with intensities as if acquired with an MPRAGE sequence. The face consists of “white matter”, the eyes of “gray matter” and the mouth of “CSF”. The data was simulated with parameters as described in the main text. In short the two leftmost phantoms were simulated with parameters as used at the FMRIB and the three rightmost with parameters as described in [33]. The middle images is simulated with homogeneous transmit and receive *B*_1_ fields, the second from right is simulated with an inhomogeneous receive *B*_1_ and the rightmost with an inhomogeneous *B*_1_ transmit field (leading to a spatially varying flip-angle).

### Registration of a single subject to the MNI152 template

A healthy 28 year-old female subject (F) was scanned on a Siemens Trio syngo system using an MPRAGE sequence with TI/TE/TR 900ms/4.53ms/2200ms respectively and an 8 degrees flip angle. The imaging matrix was 160 × 192 × 192 with an isotropic 1mm resolution and two repeats were collected. Data was collected using an 8-coil coil-array and images were reconstructed individually for each coil and combined in a sum-of-squares sense [26]. This is a representative example of a typical structural scan at the FMRIB centre. It has an easily visible bias in the anterior-posterior direction, with higher intensities towards the posterior. Most subjects will be located closer to the posterior part of the coil than to the anterior part, which is the cause of this quite typical bias. The scan was registered to the MNI152 template (the ICBM 18.5-43.5 template described in [39]) using the configuration file for T1 to MNI152 registration that is part of the FSL-package (to a final knot spacing of 10mm) and a high resolution T1 to MNI152 configuration file (to a 2mm knot spacing). Both registrations were performed using an intensity model given by Eq 16 (i.e. modelling a global nonlinear relationship between the intensities in the two images and a smoothly changing bias field) and for comparison by Eq 13 (i.e. a “standard” sum-of-squared-differences cost function). For this registration the warps were not projected back onto a diffeomorphic surface, instead the Jacobians were allowed to take negative values. This was done to demonstrate the role of intensity bias in causing non-diffeomorphic warps.

### Registration of one subject to another

A healthy 42 year-old male subject (M) was scanned using the same sequence as the female subject above, with the distinction that the matrix size was now 208 × 256 × 192. The considerably larger head of this subject (and also slight, unintentional, rotation of the head around the *z*-axis) resulted in a different bias field where the intensities were highest in the lower-left and the upper-right quadrants (as seen in an axial slice) of the brain. This was a consequence of these two parts being closest to the coil-array. F was registered to M using a configuration file for T1 to T1 registration. The registration was run to a knot spacing of 10mm and to a knot spacing of 2mm, both with intensity modelling given by Eq 16 and by Eq 13. The registration was also reversed so that M was registered to F using the same parameters as above. As in the case above warps were allowed to become non-diffeomorphic.

### Registration of atrophied subject to the MNI152 template

A healthy female 84 year-old subject was scanned on a Siemens Allegra 3T system using an MPRAGE sequence with TI/TE/TR 1000ms/3.49ms/2150ms respectively and a 7 degrees flip-angle. The imaging matrix was 144 × 214 × 200 with an isotropic resolution of 1.1mm. The subject exhibited moderate atrophy consistent with her age and the images were affected by a *B*_1−_ field with, unusually, higher intensities in the centre. The subject was registered to the MNI152 template using the FSL standard configuration for FNIRT (final knot spacing of 10mm and using Eq 16 for intensity modelling) followed by a high resolution step (to a final resolution of 2mm). The resulting warps were used to resample the subject into the MNI152 space and in addition it was inverted (see Appendix S1 for details) and the inverse warps were used to resample the MNI152 template into the subject space.

### Generation of a group template

Fifty-seven healthy volunteers with ages ranging between 20 and 62 years had structural scans as part of the fMRI protocol in studies they participated in. They were scanned on a 3T Siemens Trio using a clinical MPRAGE sequence with voxel sizes in the range 0.92 × 0.92 × 0.92mm to 1 × 1.09 × 1.09 with the majority having the size 1 × 1 × 1. Other scanning parameters (TI, TE and TR) were not completely consistent across the group, but were in the same general domain as described for the female volunteer above. All scans were affected by *B*_1−_ bias fields with gradients primarily in the *y*- and *z*-directions ranging from moderate (a factor of 1.5 across the brain) to severe (a factor of 3 across the brain). Generation of the template was started by registering all subjects to the MNI152 template after which the initial a group template was created as the average of these initial registered images. This template was refined by four more iterations of registering all subjects to the group mean followed by re-generation of the group mean. After one set of four refinements the knot spacing (warp resolution) was reduced and another four iterations were performed. This was repeated for three resolutions so that there were four refinements each with knot spacings of 10, 6 and 2mm. The full procedure was run for linear scaling (intensity model given by Eq 13) and for nonlinear scaling with bias field (Eq 16). In addition the template was created either by averaging the registered images or by averaging the registered bias corrected images so that, in total three different schemes were employed.

### Registration of surface coil Macaque data to a Macaque template

Six Macaque monkeys were scanned on a commercial full bore 3T scanner (manufacturer intentionally withheld) scanner using an MPRAGE sequence and a custom built four channel phased-array coil. The placement of the coils was restricted by the monkey being placed in the sphinx position, leading to high sensitivity (intensity) in lateral cranial parts and lower sensitivity elsewhere. The inhomogeneity of the resulting *B*_1−_ field was very severe with roughly a factor of ten between the brightest and the darkest parts of the brain and with the extracerebral tissue closest to the coils having intensities several times higher still. The inhomogeneities were severe enough to make it impossible to use FAST [40] despite it containing an explicit model for the bias field. All scans were first registered, with a 5mm knot spacing, to the MNI Macaque atlas [41] after which an initial population mean was created. The scans were re-registered to the population mean using the same registration parameters as for the registration to the MNI space. Following this five more iterations were performed of registering all scans to the population mean and re-generating the population mean. At each new iteration the knot spacing was decreased and/or the regularisation was decreased until the final iteration was run with a 1mm knot spacing. One such set of iterations was performed using Eq 13 (“standard” scaled sum-of-squared differences) and one set using Eq 16 (global nonlinear intensity mapping with a multiplicative bias field) for intensity mapping.

### Demonstration of projection onto a diffeomorphic surface

To demonstrate the effects of restricting the range of Jacobians to positive values (specifically to the range 0.01–100) we allowed registrations to run to their final resolution without any restrictions and compared that to the default strategy for FNIRT (which is to project the warps onto a diffeomorphic surface after each step in a multi-resolution–multi-regularisation execution). For this test we used the single female subject above and registered to the MNI152 template.

### Registration of NIREP data

The Non-Rigid Image Registration Evaluation Project (NIREP, [42]) consists of skull-stripped, apparently bias-corrected, high quality T1-weighted MR images from 16 healthy subjects. For each subject there are 32 manually edited gray matter, mainly cortical, regions. Each subject was registered to each other subject, starting with an affine registration using FLIRT [43] followed by a medium resolution (10mm isotropic knot spacing) nonlinear FNIRT registration with simultaneous intensity modelling (Eq 13, single scaling factor). Finally a high resolution (1mm isotropic warp resolution) registration was performed while keeping the scaling factor constant. For each registration all 32 regions were resampled and thresholded at 0.5 to yield binary label images. These were compared to the pertinent original labels for the target subject and the *Relative Overlap Metric* (or the *Jaccard coefficient*) as suggested in [42] was calculated (see definition in Appendix S1) and reported for each region. In addition we computed a modified version of the *Inverse Consistency Measure* also suggested by [42]. It was modified to use the norm, rather than the square of the norm, of the differences between the forward and backward mappings thus yielding a measure in mm that is more intuitive to interpret. The details of the modification are described in Appendix S2.

The regions appear to be very carefully defined and compared to, for example, the LPBA40 dataset the regions are more narrow, being restricted to gray matter voxels, yielding relatively large surface areas to volume ratios which should be more difficult to register and hence better discriminate between different registration algorithms.

### Comparison to results in Klein et al. (2009)

This was performed mainly for comparing how the released version of FNIRT compares to what [44] reported for an early beta version. We have relegated most of the methods and results of this to Appendix S4. In short, in [44] the information contained in the pial surface of the cortex was not used by FNIRT in contrast to all the other methods, which used this information (see footnote 4 on page 799 of the [44] paper). Therefore we re-ran the experiments in [44] on the data sets LPBA40 [45] and MGH10 [44] with identical settings except for now using the information from the pial surface of the cortex. In addition, we ran it to a 1mm knot spacing, a resolution that was not possible with the beta release that was evaluated in [44].

## Results

### Demonstration of intensity models

The results of the registrations of the simulated phantoms are summarised in table 1 and Fig 2. One of the main results that can be gathered from table 1 is that in every case the pertinent intensity model perform of the best, e.g. Eq 17 performed best for the data with inhomogeneous *B*_1+_. Another obvious result is that one pays a price by using a more complicated/permissive intensity model than is motivated given the nature of the data. Both Eqs 16 and 17 performs worse than the simpler models when there is no bias field. There is also a price in terms of computational expense: and e.g. modelling intensities using Eq 17 with a bias field knot spacing of 10mm more than doubled execution time compared to linear scaling using Eq 13. A final thing to notice is that failing to model a nonlinear mapping of intensities has very little effect compared to not modelling inhomogeneous *B*_1−_ or *B*_1+_ fields. Compare for example the performance of nonlinear scaling (Eq 15) and linear scaling (Eq 13) when registering images “acquired” with different sequence parameters (i.e. the two leftmost entries on the second row of table 1 and Fig 2). Even though the “correct” model performs slightly better, as assessed by the mean difference in displacements, the difference is very small and hardly detectable by a visual inspection of the images (though one can just about detect a difference along the left edge of the left eye). Contrast that to a comparison between the models that do not incorporate a bias field (be it *B*_1−_ or *B*_1+_) to those who do (i.e. compare the two leftmost images to the two rightmost in either of the lower two rows) where vast misregistration results from not modelling the bias field.

**Table 1.**
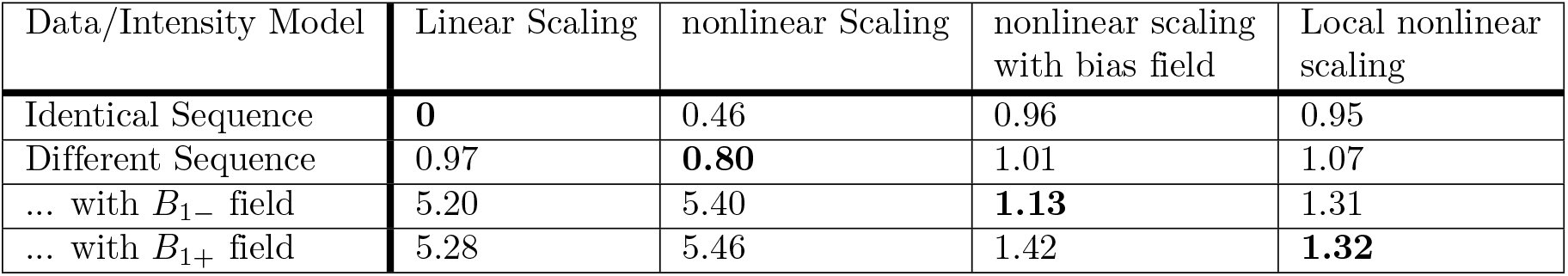
Mean difference in warp displacements (in mm) between “ideal” registration and that obtained for the pertinent combination of data and intensity model. The registration of identical intensity images without any intensity modelling was considered “ideal”. The best result for each dataset is highlighted in bold.

| Data/Intensity Model | Linear Scaling | nonlinear Scaling | nonlinear scaling with bias field | Local nonlinear scaling |
| --- | --- | --- | --- | --- |
| Identical Sequence | **0** | 0.46 | 0.96 | 0.95 |
| Different Sequence | 0.97 | **0.80** | 1.01 | 1.07 |
| ... with $B_{1-}$ field | 5.20 | 5.40 | **1.13** | 1.31 |
| ... with $B_{1+}$ field | 5.28 | 5.46 | 1.42 | **1.32** |

**Fig 2.**
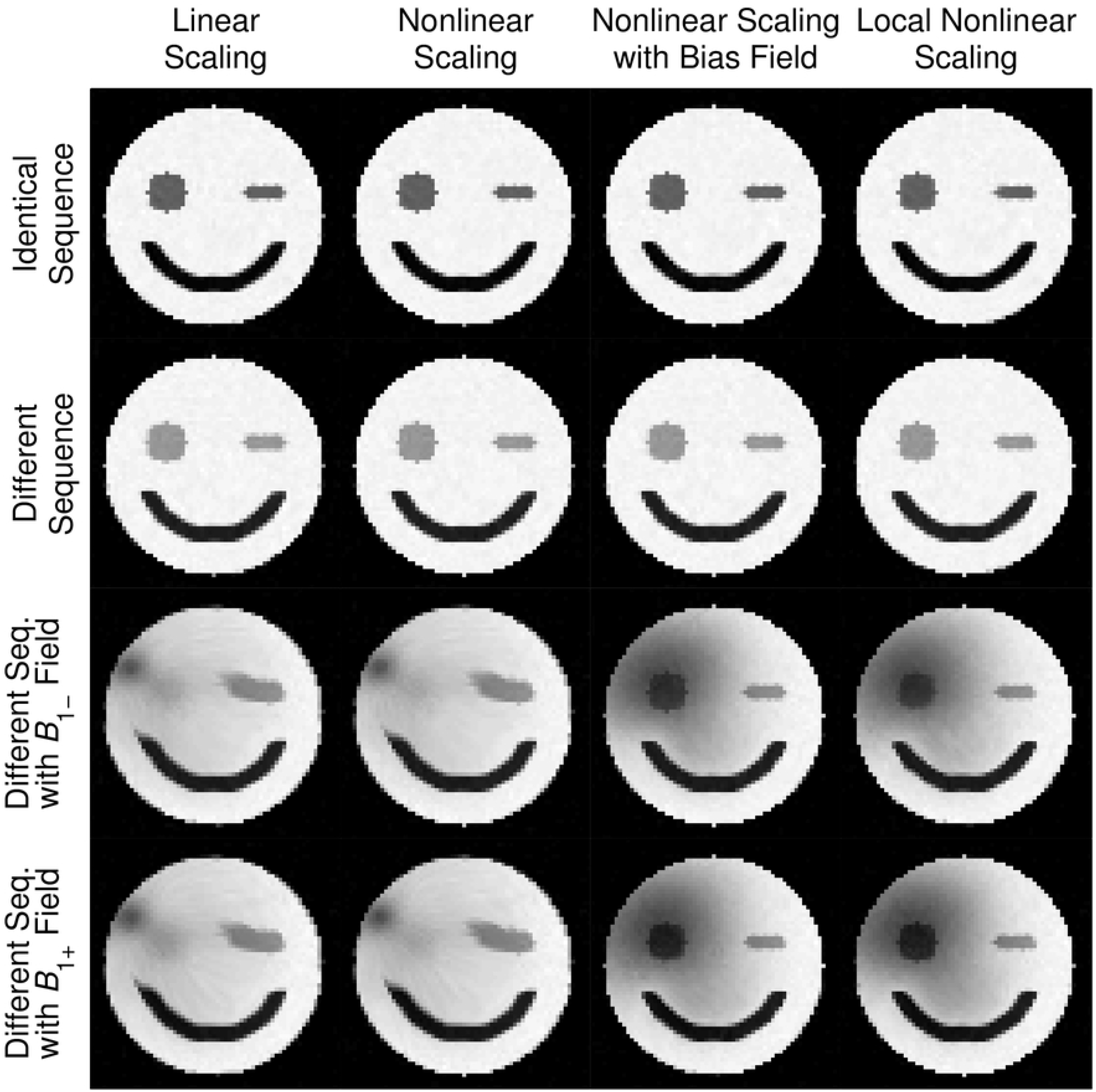
Results of registering simulated grumpy faces to a happy face. The organisation of the figure is the same as in table 1 with each row corresponding to a different simulated image and each column to a different intensity model.

### Registration of a single subject to the MNI152 template

The results, shown in Fig 3, confirm those obtained with the simulated data. The visual appearance of the registration is very good when modelling the intensities using Eq 16, but considerably poorer when using just linear scaling (Eq 13). When looking at the distribution of Jacobian determinant values one notes that “extreme” values are an order of magnitude more prevalent when not modelling intensities, indicating that intensity differences unrelated to anatomy is a crucial source of such values.

**Fig 3.**
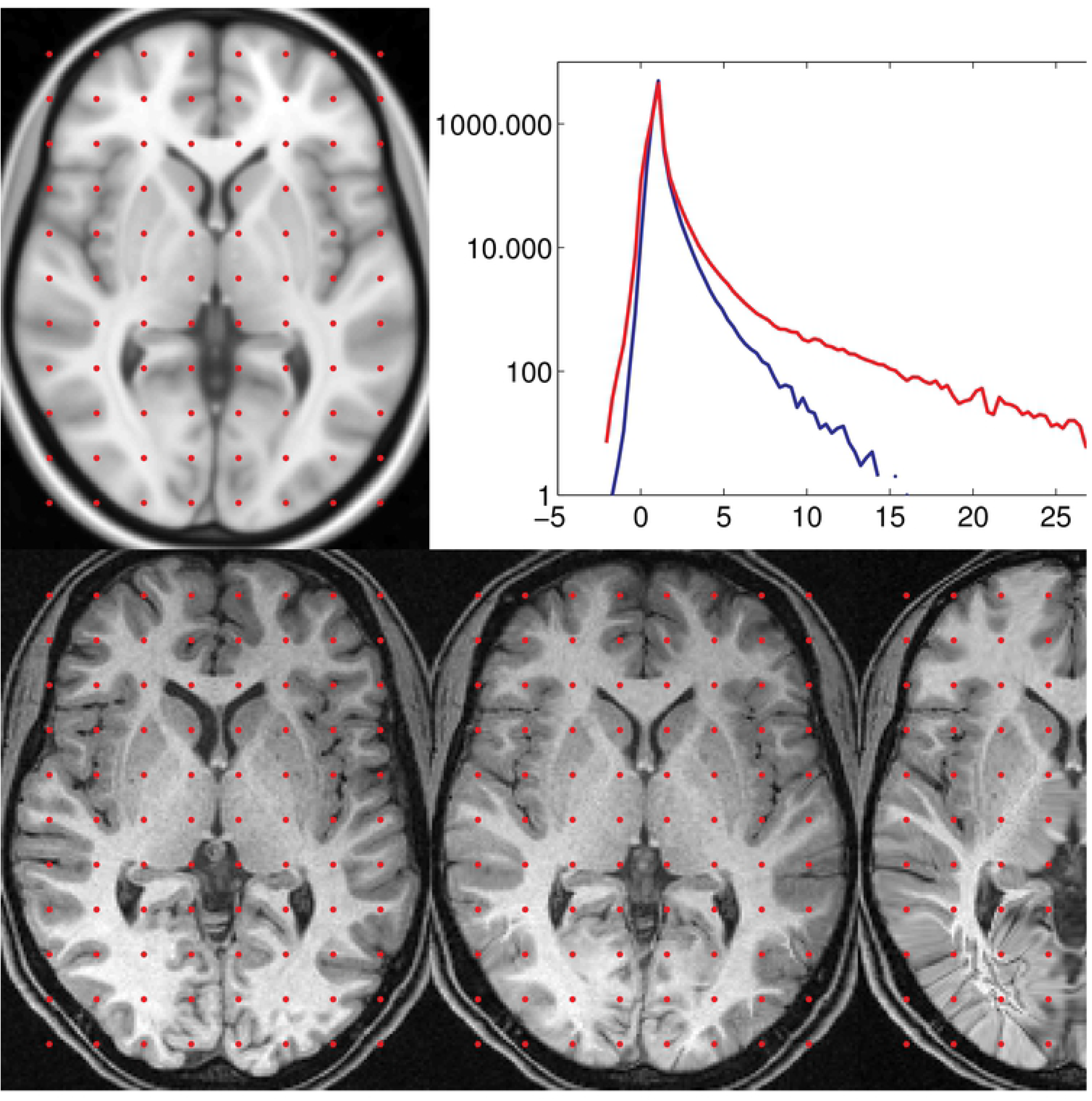
The upper left panel shows the MNI152 standard brain. The lower left panel shows the brain of a healthy female volunteer affinely registered to the MNI152 brain. Note that there is an appreciable intensity difference between the frontal and occipital parts of the brain. The lower middle panel shows the same volunteer after FNIRT registration with a 2mm knot spacing using the intensity model given by Eq 16. The image in the lower right panel has been registered to the same resolution (2mm) but using a simpler intensity model (global linear scaling). Note that the intensity model has no been used to “correct” the images, which are shown with their original intensities to demonstrate the level of inhomogeneity. The upper right panel shows the distribution of Jacobian determinants (in log-lin scale) for the intensity model given by Eqs 16 (blue) and 13 (red). N.B. that in order to demonstrate the effect of the intensity modelling the projection onto a diffeomorphic surface was turned off for this registration, and that normally there would be no Jacobian values outside the prescribed range.

### Registration of one subject to another

The subject*→*subject registration demonstrates (Fig 4) a similar behaviour with respect to intensity modelling as did the subject*→*MNI152 registration. The presence of a bias field severely disrupts the registration leading to superfluous warps when not explicitly modelled. In contrast the warps look sensible and the range of Jacobians is much reduced (reducing the number of voxels with a Jacobian determinant*≤* 0 by 50%) when modelling a nonlinear global intensity relationship and a multiplicative bias field (Eq 16).

**Fig 4.**
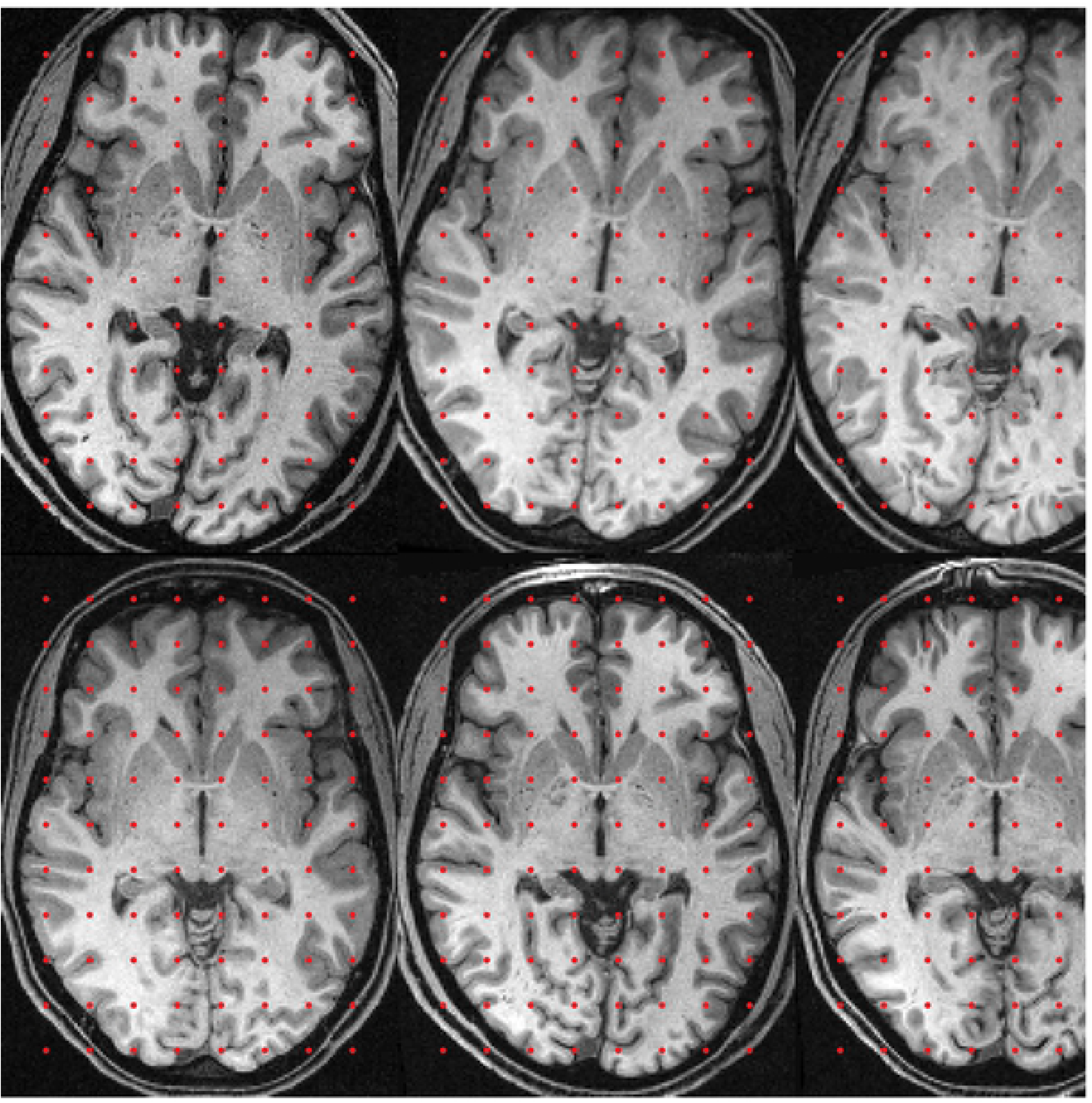
The left column shows the two target brains, the male volunteer (M) on the upper row and the female volunteer (F) on the lower. The second column in the top row shows F affinely registered to M. The third and fourth columns show F nonlinearly registered to M using Eqs 16 and 13 respectively for intensity modelling. Correspondingly the second column in the bottom row shows M affinely registered to F and columns 3 and 4 show M nonlinearly registered to F using intensity models given by Eqs 16 and 13 respectively. Note the superfluous warps in the fourth column caused by unmodelled intensity variations. All the nonlinear registrations were run to a knot spacing of 2mm.

For the subject*↔*subject registration we also demonstrate the non-symmetrical results that FNIRT yields by showing subject F transformed to the space of subject M using the warps calculated from an F*→*M registration (Fig 5, middle column) and also using the inverse of the warps obtained from an M*→*F registration (see Appendix S1 for a description of how we calculated the inverse). It is easily seen that the two solutions are slightly different, though surprisingly it is not obvious which of the two is “better”.

**Fig 5.**
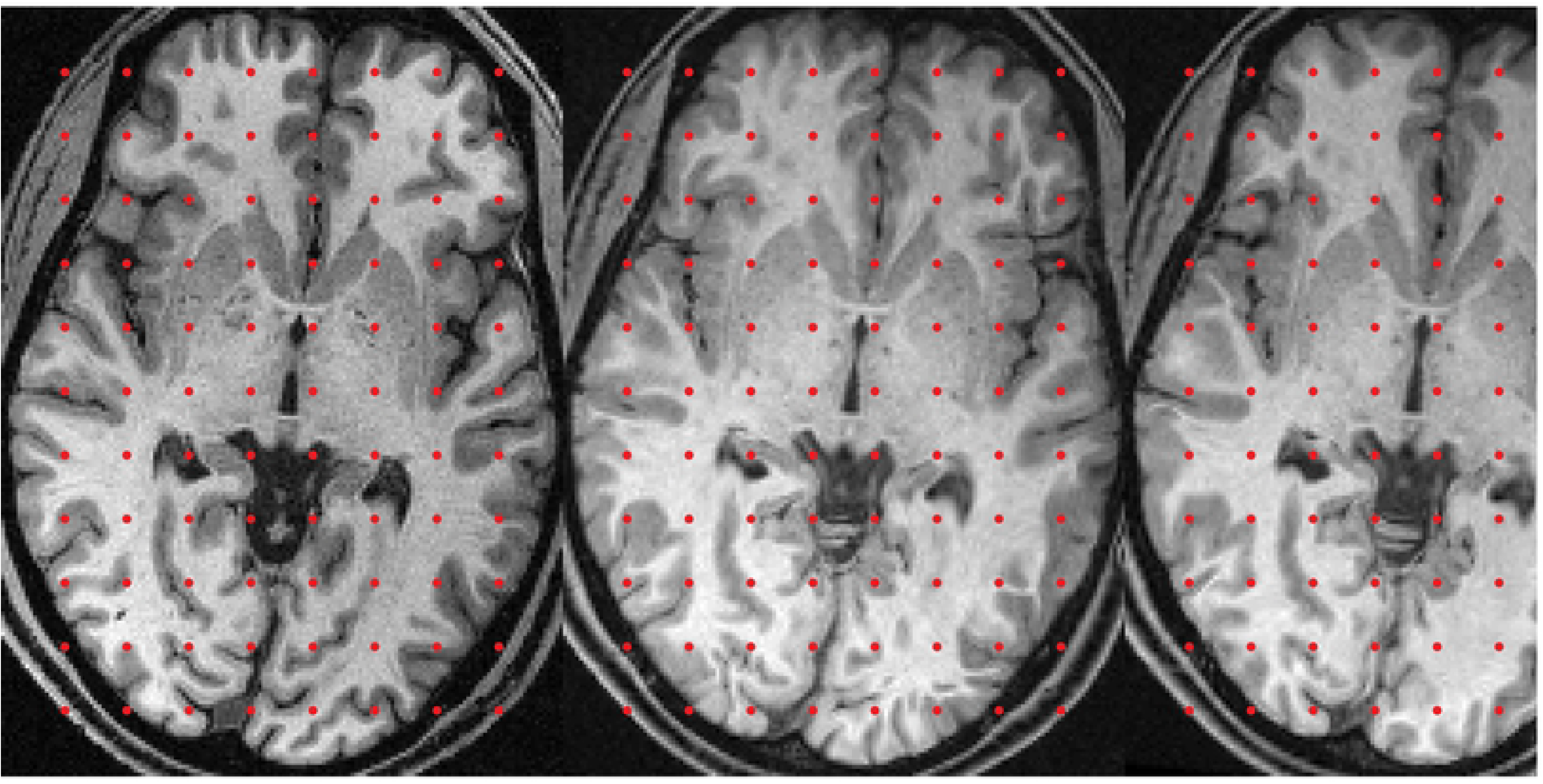
This figure is intended to show the degree of lack of inverse consistency with FNIRT. The leftmost panel shows the male volunteer (M) in native space. The middle panel shows the female volunteer (F) nonlinearly registered to M using a knot spacing of 2mm and the intensity model given by Eq 16. The rightmost panel shows F registered to M using the inverse of the warps obtained by registering M→F, i.e. the inverse of the warps that were used in the lower row of figure 4.

### Registration of an atrophied subject to the MNI152 template

Fig 6 shows the results of registering an elderly, female, moderately atrophied, subject to the MNI152 standard brain. It can be seen from the upper row that FNIRT, run to a knot spacing of 2mm, is able to register brains with this degree of atrophy with seemingly good results. The bottom row shows the MNI152 template warped into the native space of the subject using the inverse of the subject*→*MNI152 transform. The latter result was included to give some indication that even with brains this different (and the ensuing large deformations) there is a unique inverse that gives results that appear reasonable to a visual inspection. It should also be noted that this subject is in no way “selected”, but was obtained by asking around the lab for “the most atrophied subject from your study”. Our desire to stay with this “random” selection meant that it was included even though it has quite pronounced intensity changes from small vessel disease [46] which are visible periventricularly in the top middle panel and which are a possible cause of the slight, but discernible, misregistration of the white-grey matter junction of the pre-frontal cortex.

**Fig 6.**
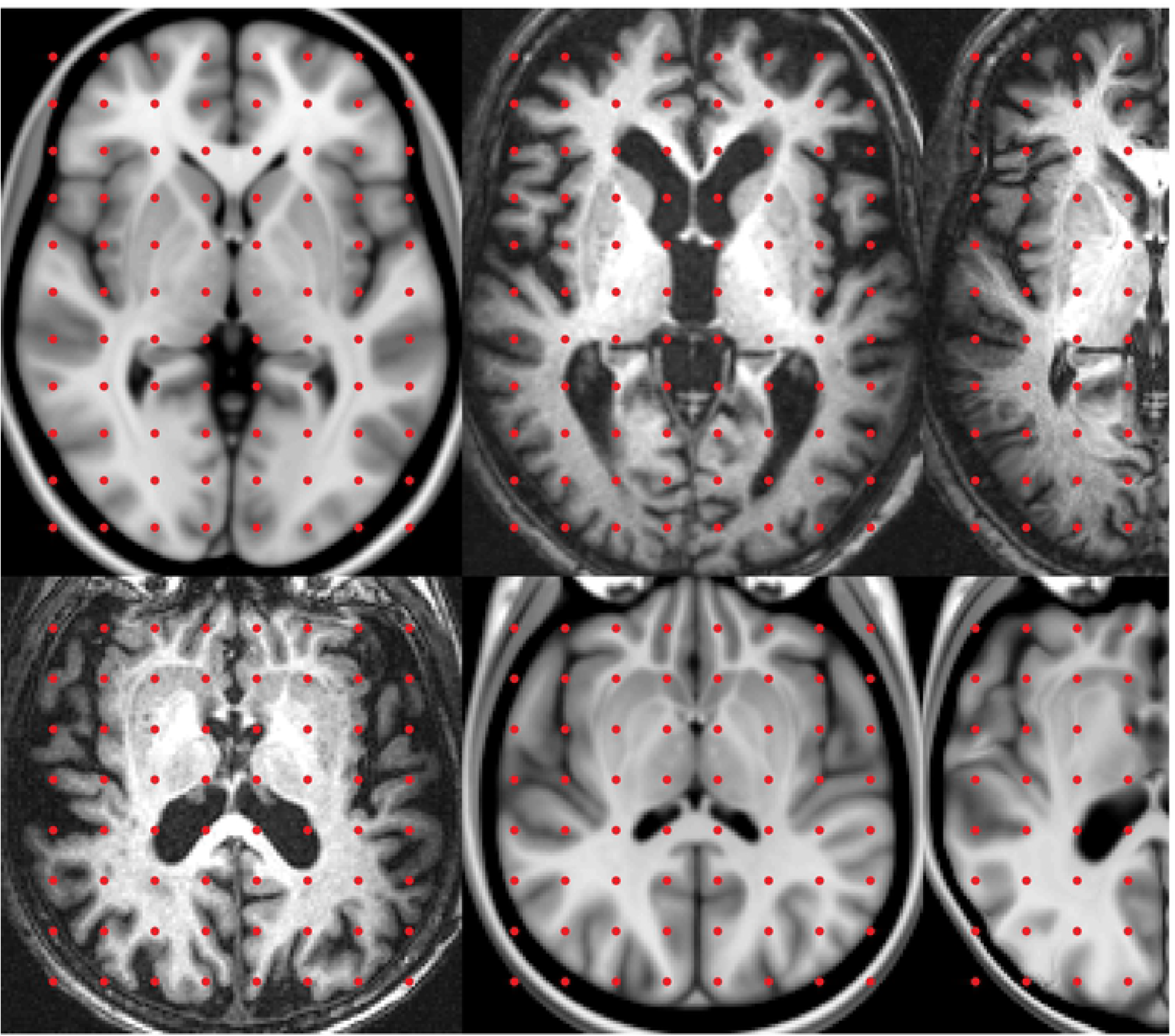
This figure shows results from registering an elderly (84 year-old), atrophied, subject to the MNI152 standard brain. The top left panel shows a slice from the MNI152 template, the top middle panel shows the same slice from the elderly subject after affine (FLIRT) registration and the top right panel after nonlinear (FNIRT) registration to a warp resolution of 2mm. The lower left panel shows the elderly subject in her native space. The lower middle and right panels show the MNI152 template transformed by the inverses of the affine and nonlinear transforms of the upper row respectively.

### Generation of a group template

The group template started drifting and did not converge when using linear scaling. In contrast it converged to a plausible group average when modelling the bias field (using Eq 16), and this was true regardless of if we used the bias corrected images for re-calculating the average or not.

An example of one slice of the group average is shown in Fig 7 together with the warped images from three of the subjects constituting this average. As can be seen, the visual agreement is very good.

**Fig 7.**
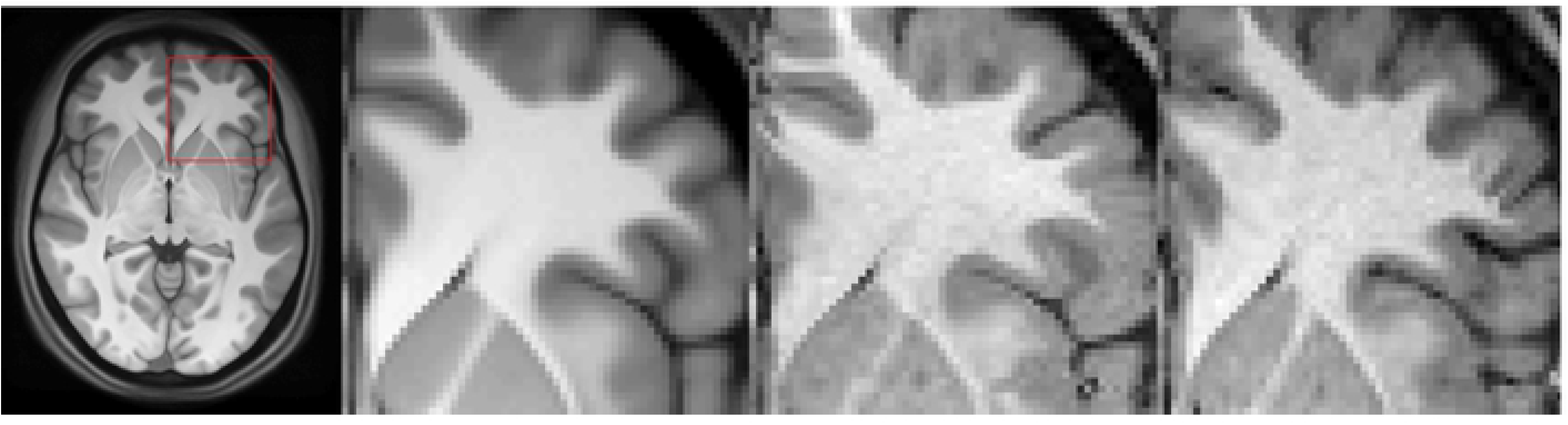
On the left is the group average with a red square indicating the part that is shown in the subsequent panels. The first of those shows again the group average, zoomed in to concentrate on a part containing ACC, Caudate, Insula and prefrontal cortex. This area was chosen as it contains the relatively “easy” structures Caudate and insular cortex, but also structures of intermediate and high difficulty such as ACC and PFC respectively. The rightmost three panels show the warped scans of three of the 57 subjects constituting the average. These were the three first subjects in alphabetical order and have not been selected as being particularly well registered. It should also be noted that the dark streak across the white matter between the Caudate and the ACC in the third subject does *not* imply a non-diffeomorphic warp, but is clearly visible in the original scan of this subject.

In Fig 8 we demonstrate how potentially interesting structure emerges from a group average created using high resolution warps even though it is not possible to discern in any of the individual images despite these being of reasonably high quality.

**Fig 8.**
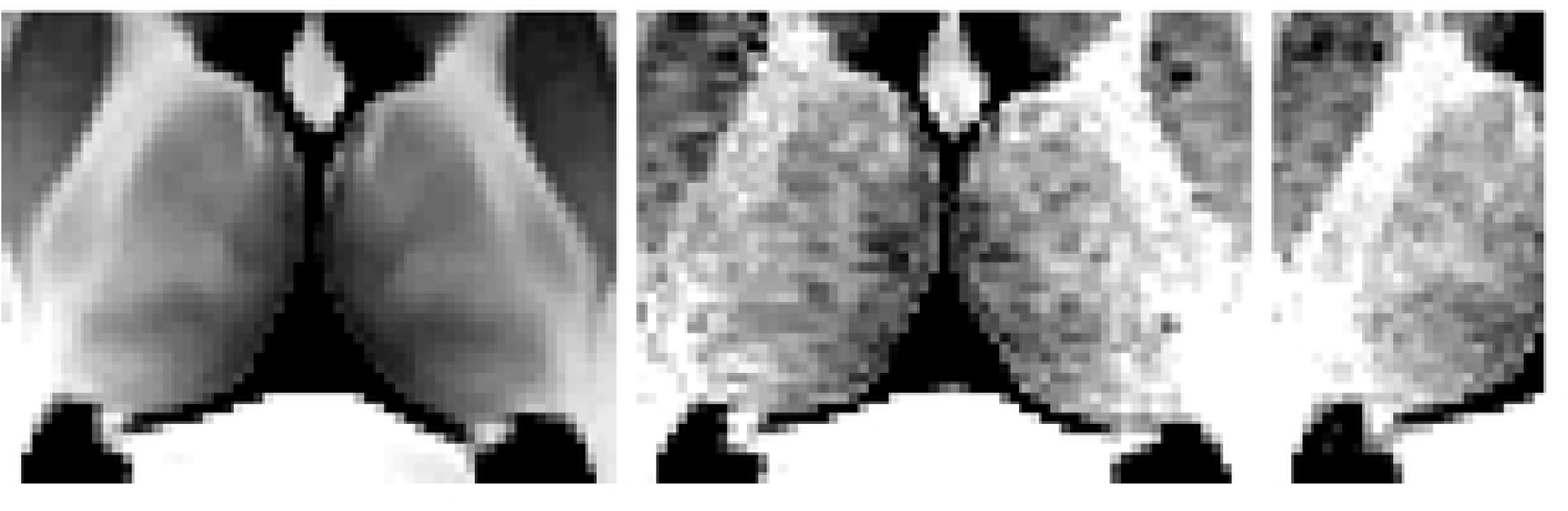
The leftmost panel shows the group average zoomed in at the region of the Thalamus and the three subsequent panels show the same area for the same three subjects as in figure 7. Note how the group map reveals a structure in the Thalamus that would not be evident by looking at an individual image. However, with knowledge of the group map this structure can actually be recognised also in the individual images, as well as the fact that this structure has been aligned across subjects.

### Registration of surface coil Macaque data to a Macaque template

When using linear scaling (Eq 13) the registered images were severely distorted and the population average did not converge. This was a consequence both of the severe *B*_1−_ field and also of the hyper-intense vessels that were present in all the scans. In contrast, when using nonlinear scaling with a multiplicative bias field (Eq 16) the registration was successful and the population mean converged to a very sharp image (Fig 9). The success of the registration can also be seen from the individual images after registration (Fig 9).

**Fig 9.**
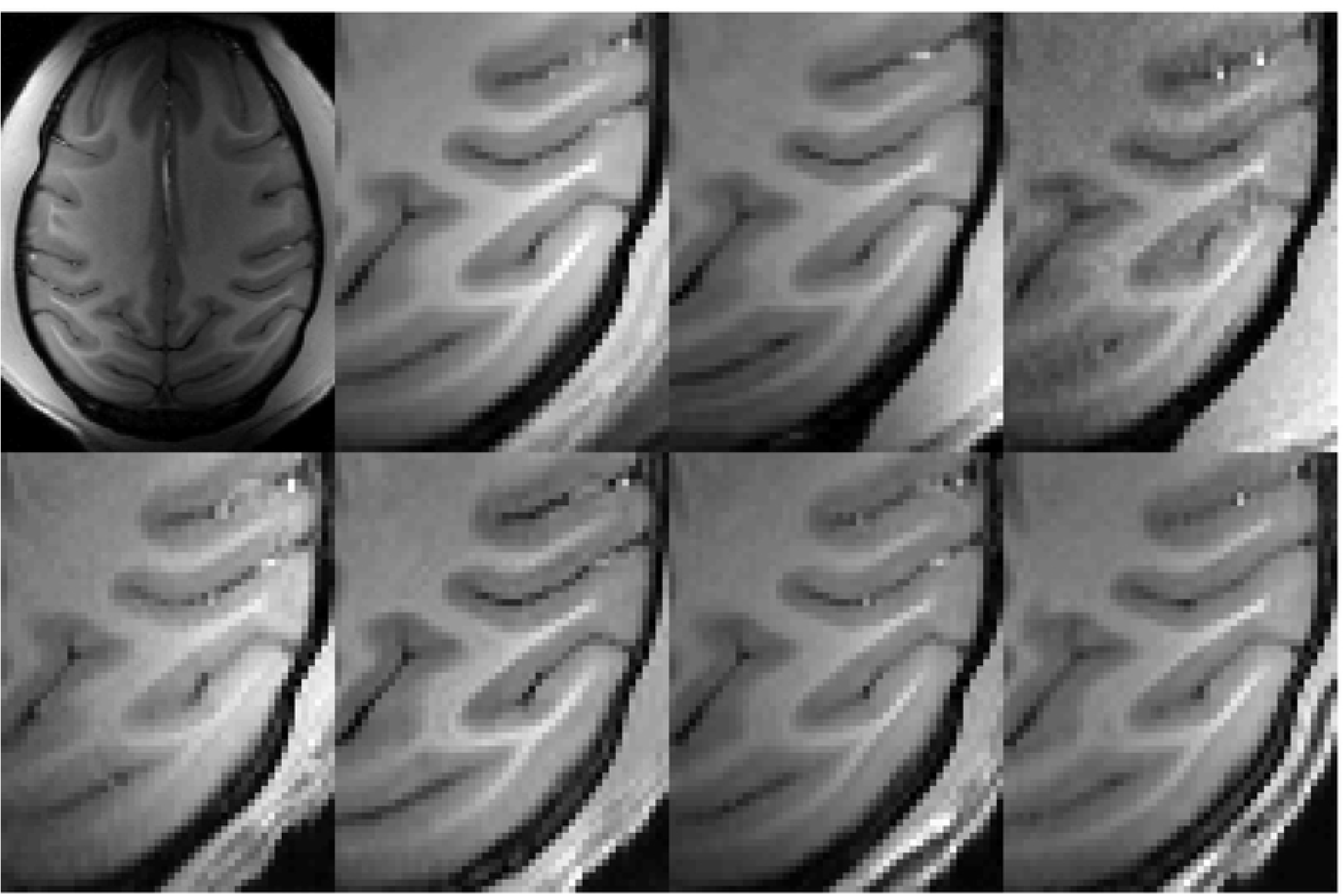
The top left panel shows an example of an axial slice of one of the Macaque scans. One can deduce the approximate placement of the four coils simply by looking at this image. The second panel from the left in the top row shows a partial axial slice of the population average of the six monkeys after registration. The remaining panels show the same area of the six individual scans after registration. It can be seen that even with a limited FOV within a slice the intensity variations are severe. In each of the displayed sections there are gray matter voxels with higher intensity than some white matter voxels, and in four of them there are gray matter voxels with intensity more than twice as high as some white matter voxels. Despite this it can be seen that the registration is good across the six subjects.

### Demonstration of projection onto a diffeomorphic manifold

A comparison of subject F registered to the MNI152 template with and without projection onto a specified Jacobian range (0.01–100 in this example) shows subtle differences. If allowed to run to 2mm knot spacing without any restrictions there were 9030 voxels with Jacobian determinants *≤* 0, while if using restrictions all Jacobians were in the range 0.01–100. Visual inspection of the registered image shows very subtle differences when restricting the range of the Jacobians compared to when not. It may seem counterintuitive that changing the displacements in excess of 9000 voxels should result in subtle, hardly noticeable by eye, changes in the registered images. However, the vast majority of non-positive Jacobians have values between −1 and 0 and typically the changes that need to be made to the displacement fields to bring these Jacobians into the required range are actually very minor.

The changes demonstrated in Fig 10 are the most obvious ones anywhere in that dataset. This is a typical finding and in no case have we come across any large or obvious changes in the registered images as a consequence of restricting the Jacobian range.

**Fig 10.**
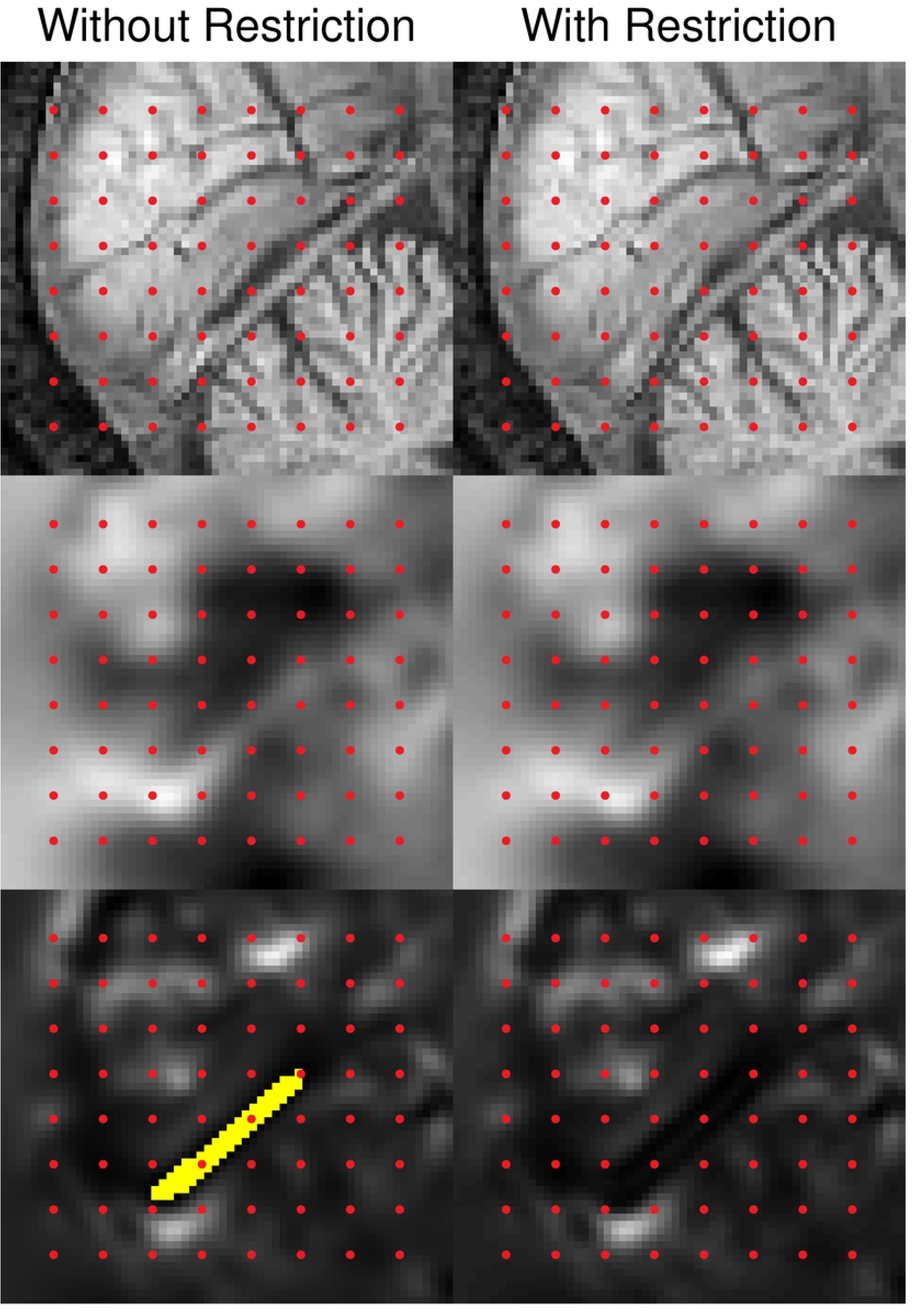
The top row shows a partial sagittal view of subject F after registration to the MNI152 template without and with restriction of the Jacobians in the left and right panels respectively. Note in particular the difference along the junction between the occipital lobe and the cerebellum. The second row shows the *z*-component of the displacement-field (scaled to −3–5mm). Note how the Jacobian restriction has widened/smoothed the relatively thin strip of positive displacements in the left panel. The third row shows the Jacobian determinants (scaled to 0–4.5). The voxels where the Jacobian was ≤ 0 are shown in yellow. Note how the Jacobian restriction has affected mainly the narrow strip of negative Jacobians in the left panel, rendering them to be within the prescribed range (0.01–100).

### Registration of NIREP data

The registration of every subject to every other subject resulted in 240 different pair-wise registrations for each of the three strategies (affine/FLIRT, FNIRT to 10mm knot spacing, FNIRT to 1mm knot spacing). An example of the overlap achieved in one of these registrations is demonstrated in Fig 11 for affine and for high resolution nonlinear registration. Fig 11 can also be used to get an impression for what (visual) overlap maps onto what Jaccard coefficient. It should be clear from the lower panel of Fig 11 that the overlap is quite good and still the Jaccard coefficient is no greater than 0.70, demonstrating the severity (and hence also efficiency) of it as a measure to gauge registration accuracy. The resulting overlaps, as assessed by the Jaccard coefficient, are shown in Fig 12. It should be noted that each box in Fig 12 represents the results of 240 registrations, each of which results in an overlap of the kind seen in Fig 11. It can be seen that, as expected, the nonlinear registration resulted in a significant improvement over the affine registration and also that the 1mm registration performed consistently better than the 10mm registration.

**Fig 11.**
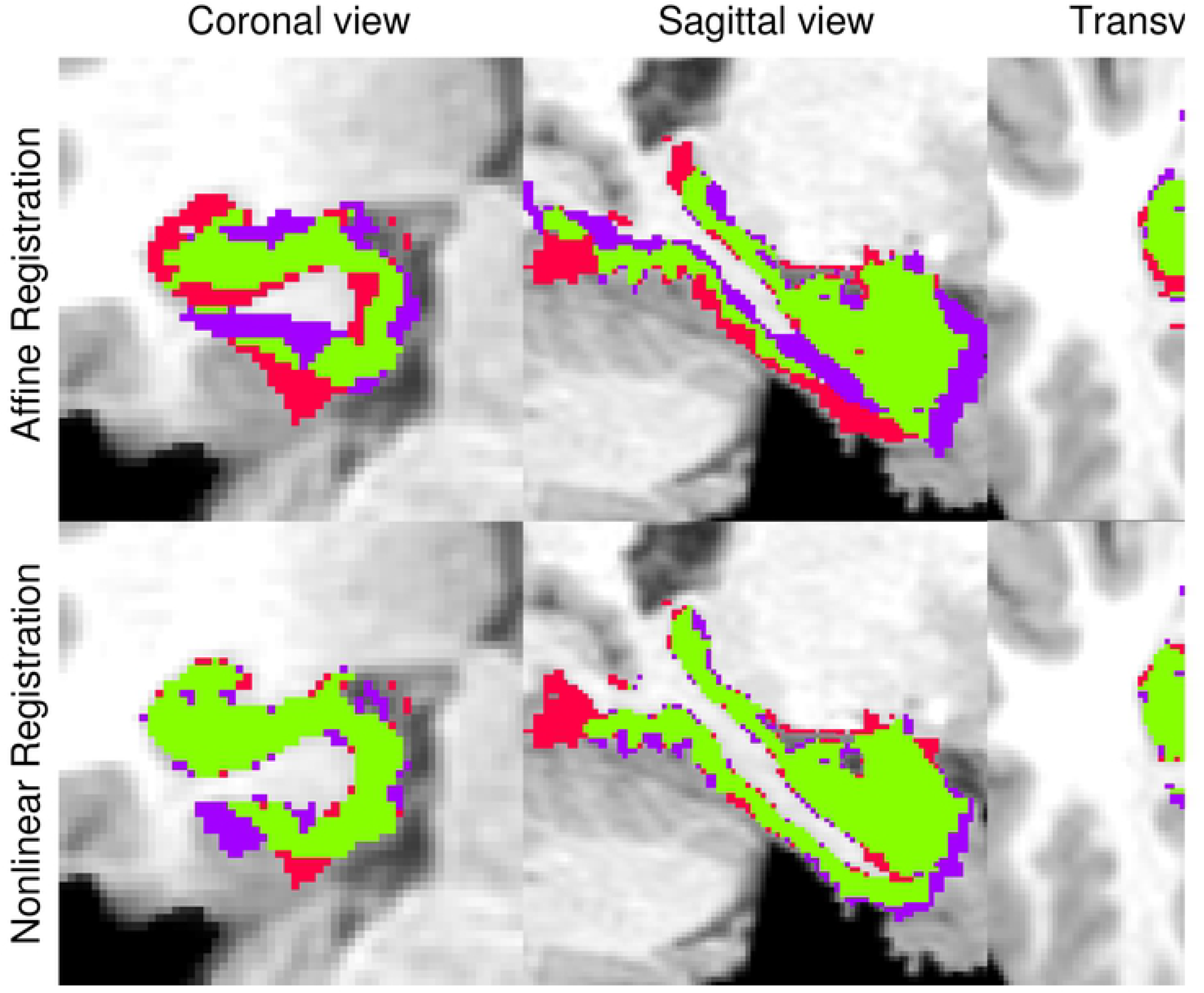
The right parahippocampal gyrus overlayed onto the scan of the target subject. The red color shows the voxels where the structure is defined in the target subject, but there is no overlap with the registered structure from the input subject. Voxels where the target is not designated as part of the structure but the registered image does designate it as belonging to the structure are shown in purple. The green voxels are those where the target and registered structure overlaps. The top row demonstrates the results for affine (FLIRT) registration and the lower row for nonlinear registration with 1mm knot spacing. For the affine case the Jaccard and Dice coefficients are 0.45 and 0.62 respectively. For the nonlinear case the corresponding numbers are 0.70 and 0.83.

**Fig 12.**
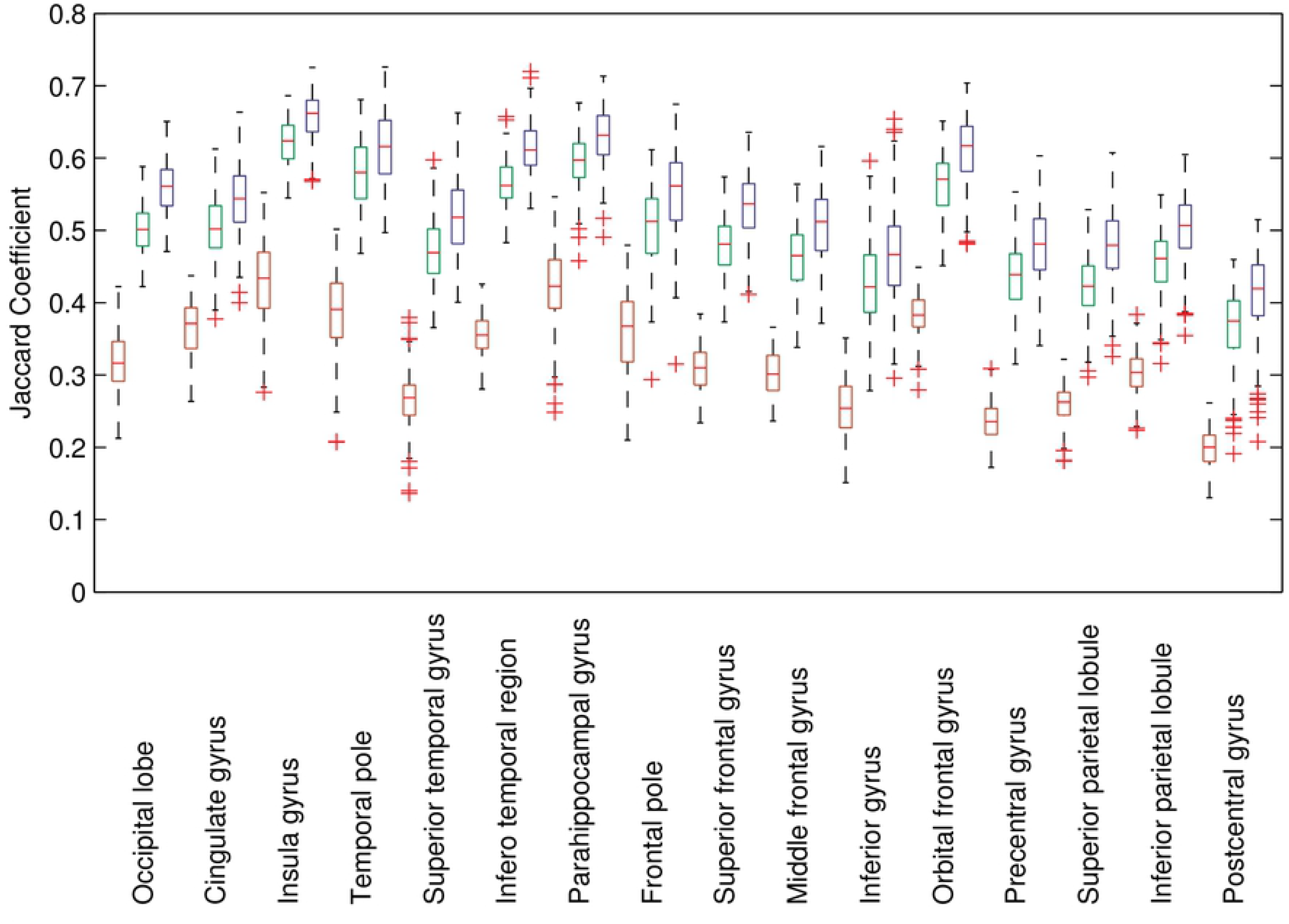
Jaccard coefficient (Relative Overlap Metric) for the 32 regions of the NIREP database, averaged across left and right sides, for affine (red) and nonlinear registration (green: 10mm knot spacing, blue: 1mm knot spacing). The line in the boxes denotes the median, the boxes extend from the 25th to 75th percentile, the whiskers indicate the total range of “non-outlier” data and outliers (approximately defined as outside ±2.7*σ*) are represented as red crosses. Each box represents 240 values (registrations). It can be seen that for all regions the nonlinear registration is a considerable improvement over the linear, ranging from 50% relative improvement for less convoluted regions with thick cortex (e.g. Insula and Parahippocampal gyri) to 100% for more difficult regions (e.g. pre- and postcentral gyri). There is also a consistent, albeit smaller, improvement when increasing the warp resolution from 10mm to 1mm.

Our Modified Inverse Consistency Error (*MICE*) is shown in Fig 13. As expected the consistency is significantly worse for the nonlinear registration than for the affine registration. It should be kept in mind though that performing no registration at all would result in a perfect consistency. It is interesting to note that the *MICE* actually decreases (for most regions) when going from 10 to 1mm knot spacing. This, together with the higher overlap scores, is a strong indication that the increased resolution really results in a more “correct” registration. The voxel wise *MICE* averaged across all registrations is shown in Fig 14. Note how the *MICE* is largest in the parietal cortex. This is consistent with the subjective impression that the variability in location (and even existence) of sulci is greater here than in the rest of the brain.

**Fig 13.**
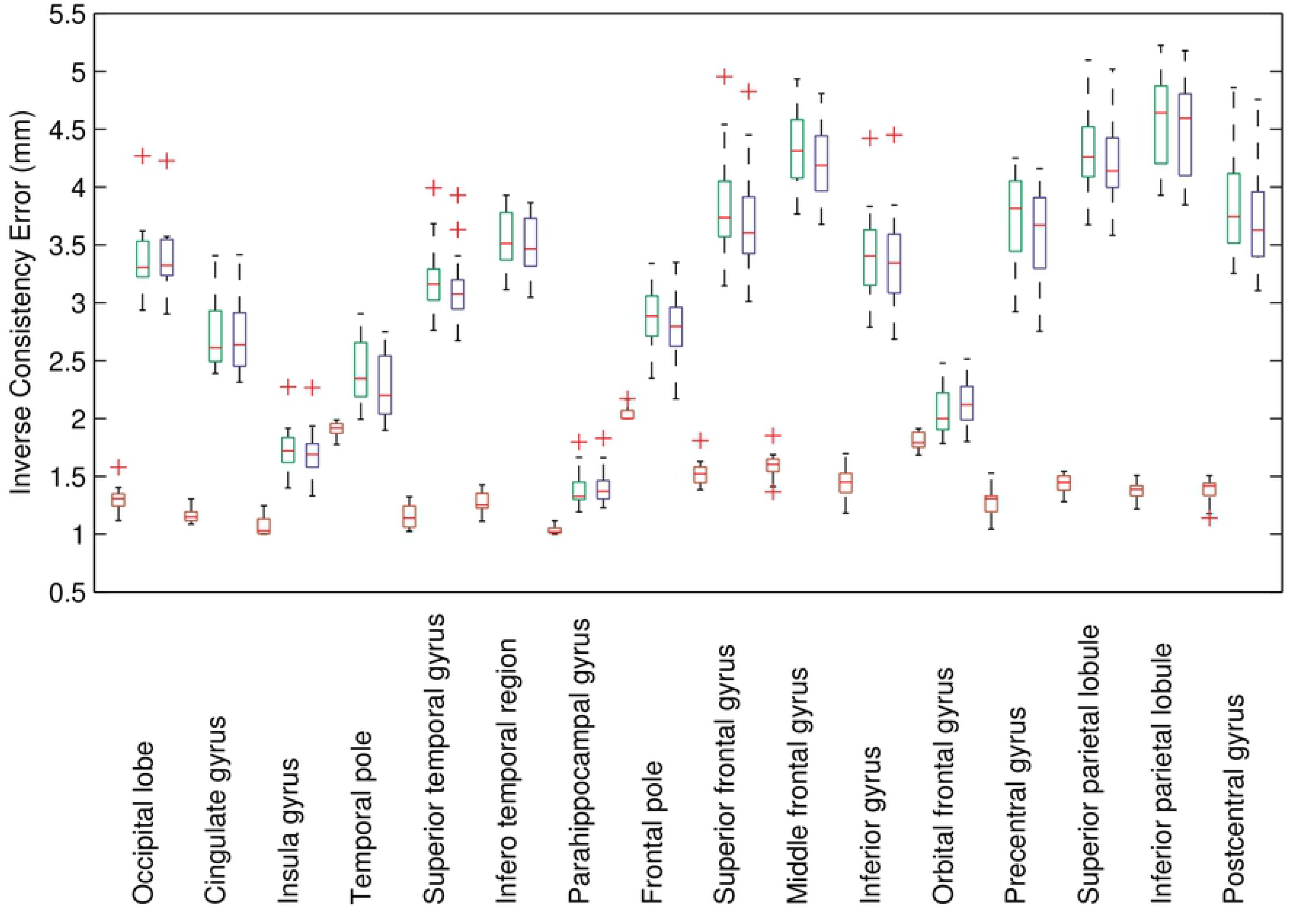
Modified Inverse Consistency Error (*MICE*) averaged within the 32 regions and across left and right sides for affine (red) and nonlinear registration with 10 (green) and 1mm (blue) knot spacing. The line in the boxes denotes the median, the boxes extend from the 25th to 75th percentile, the whiskers indicate the total range of “non-outlier” data and outliers (approximately defined as outside ±2.7*σ*) are represented as red crosses. Each box represents 18 values (subjects), each of those values being an average across 17 registrations. The pattern here is partly the opposite from for the overlap criteria, with the linear registration having considerably smaller error for all regions. It can also be seen that for the “easier” regions such as the Insula and Parahippocampal regions both the errors and the difference between linear and nonlinear registration is small, while it is large for “difficult” regions. However, the pattern is not “just” the opposite of that for the overlap criteria in that a “difficult” area such as the pre- or post-central gyrus has a smaller *MICE* than e.g. the Superior or Inferior parietal lobules. Note also that the *MICE* tends to be smaller for registration with 1mm knot spacing than with 10mm knot spacing.

**Fig 14.**
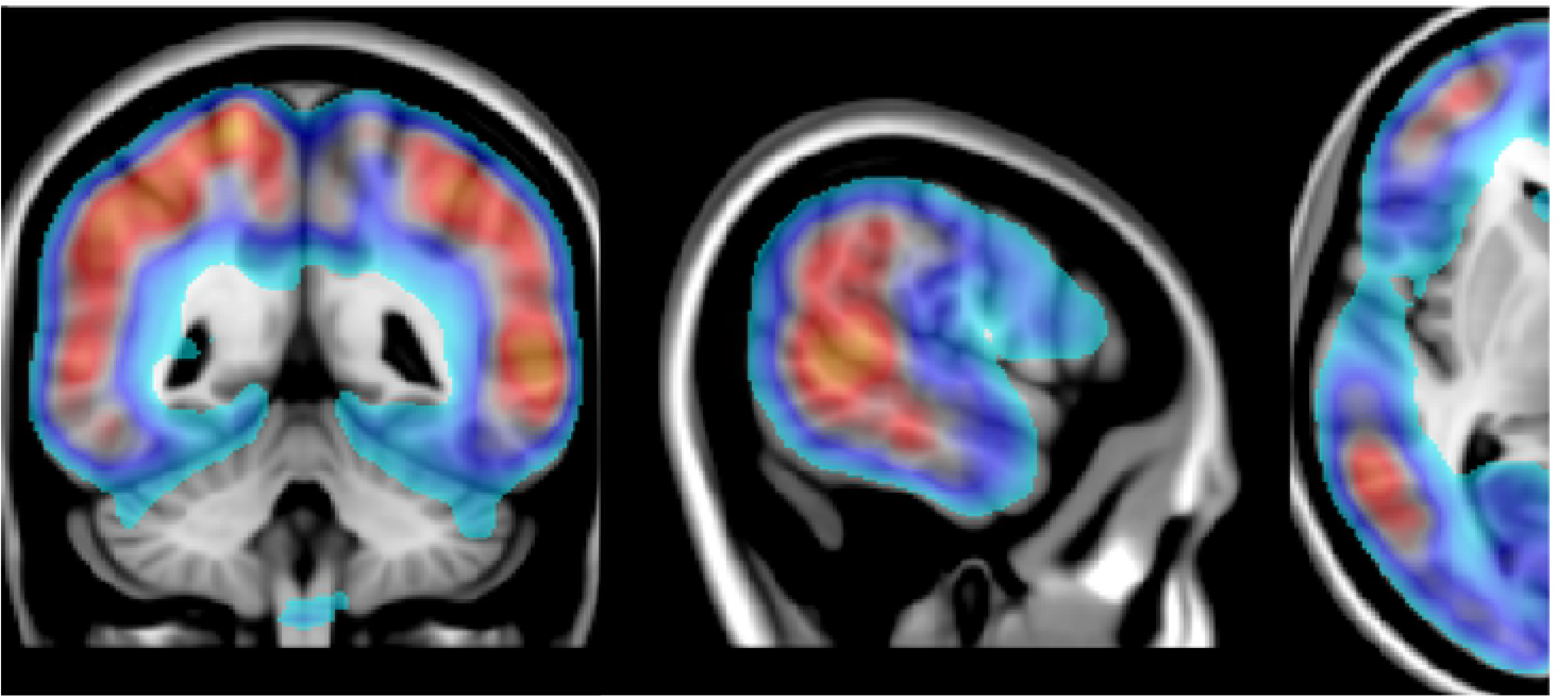
Map of Modified Inverse Consistency Error (*MICE*), as defined in Appendix S2, averaged over all 16 subjects of the NIREP database. The *MICE* was calculated for each subject (in the space of that subject) based on forward and backward registrations (to a knot-spacing of 1mm) to the other 15 subjects. The subject specific *MICE* maps were then registered to the MNI space using an affine transform and averaged across all subjects. It has been thresholded so that *MICE* <2mm is not color coded and values for 2mm < *MICE* < 6mm are coded from light blue to bright orange. The maps have been overlayed on the 1mm MNI152 template to aid anatomical orientation. Note how the consistency errors are large in the parietal and prefrontal cortices, smaller in primary sensory and visual cortices and smallest in central structures.

### Comparison to results in Klein el at. (2009)

The results from the re-analysis of the LPBA40 and MGH10 data confirmed that FNIRT performed significantly better when using the information from the outer cortical surface. It also showed that running FNIRT to a knot spacing of 1mm further improved the results. The exact ranking of the different methods depended a little on what overlap measure was used and on the details of how the ranking was performed.

For the LPBA40 data the top methods were ranked FNIRT-HR, ART/SyN, FNIRT-LR/JRD-fluid, where for example “ART/SyN” indicates that ART and SyN would alternate for second and third rank depending on overlap measure and where ART, SyN and JRD-fluid refer to the metods described in [47], [29] and [48] respectively. FNIRT-HR and FNIRT-LR refer to FNIRT run to a knot spacing of 2mm and 10mm respectively. For the MGH10 data the top methods were ranked SyN/FNIRT-HR and ART/FNIRT-LR with no other method achieving a rank among the top four.

The full set of results is presented in Appendix S4.

## Discussion

### Modelling intensities

The novel aspect of the present work is the simultaneous modelling of image intensities and geometric warps. As we have demonstrated with both simulations and example data the impact on the estimated warps from unmodelled intensity variations can be substantial. The data presented in this paper and our experiences of using FNIRT indicates that it is primarily local variations from inhomogeneous *B*_1−_ and *B*_1+_ fields that is an issue. Global nonlinearities in intensity matching, such as differences in degree of T1-weighting, seem to have a considerably smaller impact. It is not yet clear if it is necessary to make the distinction between the effects of a *B*_1−_ and a *B*_1+_ field (i.e. if it is generally useful to use Eq 17 instead of Eq 16) and as of present the recommendation is to use Eq 16. More experience of data from very high field scanners (where *B*_1+_ inhomogeneity is a greater problem) will allow us to investigate this in more detail in the future.

Our evaluations indicate that the simultaneous modelling renders the method very robust to the severe bias fields resulting from “sum-of-squares” reconstruction of data acquired with coil arrays (as opposed to bird cage coils that result in more homogeneous images). This is an increasingly important property of a registration algorithm as coil arrays with increasing number of coils are becoming more common even in clinical settings. An alternative to explicit modelling of the intensity mapping would be to use pre-processing such as histogram equalisation [49] and bias correction (e.g. [50]). We have opted for a different strategy mainly because we believe the two processes jointly inform each other. If we know the true (global and local) mapping of intensities we can perform an accurate registration. Correspondingly if we know the true spatial mapping it would be easy to find the intensity mapping. Hence none of the problems can be said to precede the other in any meaningful sense and they benefit from being addressed simultaneously.

### Comparison to other methods

The published strategy that is most similar to ours is the combined segmentation-registration framework in SPM [32]. Similar to our model, theirs includes an explicit representation and modelling of a bias field. In the segmentation model each brain tissue class (gray and white matter and CSF), and some classes intended to represent extracerebral tissue, is represented by a mean and dispersion. The registration is achieved by warping population based tissue priors to the subject’s data so as to maximise the posterior probability. This gives it even greater flexibility than our polynomial model in how intensities are globally matched between two images. Any image, regardless of contrast, can be registered to a set of prior probability maps. On the other hand the segmentation does potentially discard useful information from differential intensities within a given tissue class that FNIRT is able to utilize. For example subcortical gray matter structures will have a different T1 compared to cortical gray matter [38], and the central sulcus has been implicated as having a different T1 than other cortical areas [51]. Additionally, some subcortical areas have structure that can potentially be used to guide registration (see figure 8) but that is unused in a joint segmentation-registration framework. It will also be a disadvantage to have to perform two registrations to a set of tissue priors when registering two scans of the same subject in longitudinal studies. A consequence of the choice of basis-function in [32] is that they are limited to a rather crude warp resolution (20mm), but when a higher resolution is called for they augment that with a DARTEL registration [18] of gray and white matter segments. This is similar to our strategy of fixing the intensity model for the final resolution steps when estimating the warps.

A different strategy is represented by methods such as those by [30] and [29] where a “global” cost function has been rendered local by sampling it locally. Both these groups show promising results, and the latter has additionally performed very well in a recent comparison of nonlinear image registration methods (see [44] and section) though the specific problem of locally varying intensities (bias field) was not addressed in that study.

### Quantitative evaluation: The NIREP data

The main quantitative evaluation was performed on the NIREP data [42], though we did also use the LPBA40 [45] and MGH10 [44] data. The regions constituting the NIREP database appear (to a visual inspection) to be very carefully defined and follow both the pial cortical and the white matter surface faithfully. It is our subjective impression that it is superior to other databases of the same type (e.g. LPBA40 [45] or MGH10 [44]) which is the reason it was our main choice for the quantitative evaluation. There is a limited amount of reported results using the NIREP database, but our results compare favourably to those we have found (i.e. [52]). In the main body of the paper we graphically present some results for the Jaccard coefficient (Relative Overlap in [42]) and for our modified *ICE* index. In the Appendix S3 we also present tabulated values for overlaps and consistency error that might be useful for comparison to future studies/methods.

### Limitations of datasets for quantitative evaluation

These datasets (NIREP, LPBA40 and MGH10) had all been highly “massaged” and were all bias field corrected (though this is not explicitly stated for the NIREP data) and skull-stripped. This means that data that is used for evaluation of registration methods is not necessarily representative of the kind of data that a widely used registration software tool might expect to encounter. In particular, the manual skull-stripping introduces a very sharp (and supposedly accurate) edge in the images that will be driving registrations. Non skull-stripped data makes the problem considerably more difficult for two reasons. Firstly, T1-sequences used for brain imaging are expected to give a high gray/white matter contrast and the parameters are typically tuned for that. Considerably less attention is given to the intensities in extra-cerebral tissue such as the meninges and the spongy bone of the skull. The consequence of that is that the intensity in these parts can vary considerably between scans from different scanners and/or sequences. This is especially problematic for the meninges since that intensity is right next to the pial surface of the brain. In different T1-weighted scans we have seen intensities in this tissue ranging from much lower than in gray matter to roughly equal to gray matter to higher than gray matter. This causes problems when attempting to register two scans with different intensities in the meninges and it is not unusual to see the edge of the meninges in one scan registered to the pial cortical surface in the other. Secondly, the sharp edge coinciding with the pial surface of the cortex in skull-stripped data may or may not be present in non-stripped data depending on the intensities in the meninges. For cases when that edge is weak or absent the alignment of the cortical surface is likely to be worse.

This means that quantitative results from data like these will reflect a best case scenario and that both overlap and inverse consistency are worse in the “normal” clinical or scientific laboratory setting. It also means that when comparing methods, that aspect of how they deal with these kind of real world problems is not addressed.

### What resolution warps should FNIRT be run at?

Despite the concerns expressed above, one thing was learned from the quantitative evaluation. That is, what resolution warps are supported by scalar structural data, i.e. what resolution warps is meaningful to try to estimate given typical data. The results from the validations yield slightly different answers, where the LPBA40 and MGH10 datasets indicate that there is little gain in going from 10mm to 1mm resolution whereas for the NIREP set there seems to be a significant advantage. Our interpretation is that the data itself (the MR scans) are of similar quality in the three sets but that the regions have been more carefully delineated in the NIREP set, and that because of this it is possible to validate the results to a higher warp resolution than 10mm. For the two other datasets (LPBA40 and MGH10) the regions appear to be less carefully defined and therefore less suitable for demonstrating the advantage of high resolution warps. Hence we believe that, if the application motivates it, it is meaningful to run FNIRT to a higher resolution than 10mm.

The motivation for using lower resolution is often to decrease execution time [53] or memory requirements ([11] and [32]). If that is not an issue we have found it advantageous to run the warps to a high resolution and regulate the smoothness by *λ* (as defined in Eq 20). To enforce smoothness by setting a fixed knot spacing greater than the voxel size leads to unwanted consequences since not all warps can be represented. This often leads to “compromises” where the images appear identical, the sum-of-squared differences are (close to) identical but where the warps represented by the larger knot spacing are non-diffeomorphic while the warps generated from a smaller knot spacing are diffeomorphic. There seems to exist a misunderstanding that representing the warps by a set of smooth basis-functions helps protect against negative Jacobians, and that a larger knot spacing (or lower cut-off frequency) offers a better protection. This is not the case and a larger knot spacing simply means that the regions containing non-diffeomorphic warps are bigger and that there is smooth transition from the diffeomorphic to the non-diffeomorphic regions.

Hence our recommendation would be that, as long as execution time is not an issue, one runs FNIRT to the highest resolution supported by the reference image and then uses the regularisation to tailor the smoothness of the warps to the particular application (it may, for example, be advantageous to use a greater smoothness when registering fMRI group studies).

### Diffeomorphic by construction: are there any practical advantages?

For some years there has been a tendency towards methods that are “diffeomorphic by construction”, i.e. methods that update the warps in such a way that if the update itself is diffeomorphic then the resulting new warps are too. This means that (e.g. by performing enough iterations) one can realise displacements (warps) of an arbitrary magnitude while still being (almost) guaranteed diffeomorphic warps as long as each update is sufficiently small/regularised. In contrast, our method uses what is known as a small deformation framework, a framework that has been perceived as not being able to produce large deformations without incurring a non-invertable field. Another difference between the two strategies is that in “diffeomorphic by construction” methods regularisation is typically applied to each update instead of to the resulting field. This means that one does not “build up tensions” in the warps which can therefore grow to any magnitude without incurring any cost. In contrast, small deformation methods typically regularise the warps themselves, using that as a means to prevent non-invertible warps but at the same time limiting the warps that can be accommodated.

Despite these purported limitations we opted to develop our method within a small deformation framework. The reason for this was our desire to include a simultaneous intensity model that would allow us to address imperfections in real data such as bias fields and the difficulties in estimation the gradient and Hessian of such a model within a “diffeomorphic by construction” framework. So, in order to attempt to overcome the disadvantages of the small deformation model we implemented a multi-resolution, multi-regularisation scheme with projection onto a diffeomorphic warp after every few iterations.

In the present paper we demonstrate that this has yielded a method that achieves equally good/better overlap of manually delineated regions compared to other methods (based on the comparisons with the results in [44]) and which still only produces diffeomorphic transformations. Given this, it is unclear to us what advantages the “diffeomorphic by construction” framework has to offer (besides mathematical elegance) by comparison with the projection framework used here.

### More difficult applications

All the data which we have used for “proper” numerical validation and comparison (NIREP, LPBA40 and MGH10) have consisted of scans of young to middle-aged healthy volunteers. Our results indicate that for registration within these kinds of populations a small deformation framework along with careful implementation and projection onto a diffeomorphic manifold is sufficient to obtain good results. It is possible that for more difficult applications, e.g. for registering severely atrophied to healthy brains, or monkeys to humans [16], diffeomorphic by construction algorithms may have an advantage. This is however something that remains to be demonstrated and, as demonstrated in figure 6, we have used FNIRT for registration of atrophied subjects with seemingly good results while still producing diffeomorphic mappings. The examples that are often used to demonstrate the advantages of “diffeomorphic by construction” methods, such as the patch-to-a-C example [9] or moving a control point through a line/plane defined by other control points [16], are somewhat contrived and not necessarily representative of anything one would ever encounter while registering scans of brains.

### Inverting and composing warps

When calculating a diffeomorphic warp in an Eulerian framework it is trivial to calculate also its inverse and, possibly more important, one can also calculate the gradient and Hessian of the cost function for both forward and backward warps [18]. This means that it is feasible to estimate *exactly* inverse consistent warps (e.g. [18] and [29]), something that would not be possible in a small deformation framework. This is a potentially important point since the constraint imposed by inverse consistency when warping one scan to another is a highly reasonable and unequivocal one, unlike, for example, the regularisation model that is arbitrary (i.e. membrane energy, bending energy or linear elastic energy) and chosen more for computational convenience than for empirical reasons.

It is, however, feasible to calculate a good approximation the inverse of warps obtained with a small deformation framework [54], and it can be done in a few tens of seconds (see Appendix S1). Hence, the implicit assumption in [18] that the inverse warps in a small deformation setting is typically approximated by **x** − **d**(**x**) is neither necessary nor common.

### Comparison to the results reported by Klein et al. (2009)

Our re-evaluation of the data used in [44] confirmed our hypothesis that FNIRT would perform better if using the information in the pial edge of the cortex. All the evaluations in [44] were performed on skull stripped data. Skull stripping introduces an artificial sharp edge of non-zero to zero voxels where the skull stripping algorithm/observer locates the cut, which means that the accuracy of a subsequent registration will depend crucially on the accuracy of the skull stripping. For this reason we usually recommend that FNIRT is run on non-skull stripped (original) data unless one has a very good reason to believe that the skull stripping has performed perfectly. If, for some reason, only skull stripped data is available we usually recommend that one runs FNIRT with a switch that tells it to regard zero values as missing data, thus ignoring the artificial edge (as long as one is not 100% confident in the accuracy of the stripping). For these reasons we initially recommended these settings (ignoring zeros) for the evaluations performed by [44], unlike all the other methods that used the artificial edge.

What we did not consider at the time was that it didn’t matter if the brains were poorly extracted or not since the extraction was based on the union of all manually labeled regions. This means that if one wants to perform well in terms of aligning these regions one should definitely use the information in the artificial edge since it pertains directly to the measure that is used for the evaluation. Unfortunately this creates a strong dependence between the “data” (the skull stripped scans) and the “ground truth” (the manually defined labels). These should of course ideally be independent. Paradoxically this means that the it would be more “valuable” to use the information contained in a brain*↔*zero junction than to use any other internal junction regardless of how “objectively wrong” the former is.

Despite these reservations regarding the evaluation of registration algorithms on data of this kind we find it encouraging that FNIRT performed so well in comparison to other methods. Both the data (LPBA40 and MGH10 (https://www.synapse.org/#!Synapse:syn3207203)) and the software (https://fsl.fmrib.ox.ac.uk/fsl/fslwiki/FslInstallation) are publically available for anyone wishing to confirm our findings.

It should be noted that FNIRT was not the only method that was a beta release at the time and it is conceivable that some of the other methods would also perform better if their finished, released, versions were used (see e.g. [35] for a re-evaluation of DARTEL).

## Conclusion

We have described the implementation of a method/framework for small-displacement nonlinear registration of brain MR images. The results look promising and on “ideal” data compare favorably to the methods included in [44]. However, we believe that the main advantage of the presented method is on the “imperfect” data commonly encountered in a clinical or neuroscience setting, especially with respect to bias fields commonly seen in modern coil array acquisitions.

Future work will focus on a multivariate framework where the intensity modelling is extended to a general model-driven transfer function.

## Supporting information

**Appendix S1 Additional information about the method.** Additional information about the details of the implementation and a description of a fast method for calculating the inverse of a warp.

**Appendix S2 Evaluation criteria.** Definition of the metrics used in the quantitative evaluation.

**Appendix S3 Additional results from the NIREP evaluation.** Tables and figures with results from the NIREP evaluation not included in the main text.

**Appendix S4 Comparison to results from Klein et al. (2009).** Results from re-analysis of two of the datasets used in [44] (the MGH10 and LPB40 data sets) using different parameters for FNIRT.

